# The IFIH1-A946T risk variant promotes diabetes in a sex-dependent manner

**DOI:** 10.1101/2024.01.20.576482

**Authors:** Amanda J. Stock, Pierina Gonzalez-Paredes, Luciana Previato de Almeida, Stanley D. Kosanke, Srinivaas Chetlur, Hannah Budde, Paul Wakenight, Theresa A. Zwingman, Aaron B. Rosen, Eric Allenspach, Kathleen J. Millen, Jane H. Buckner, David J. Rawlings, Jacquelyn A. Gorman

**Affiliations:** Oklahoma Medical Research Foundation, Arthritis & Clinical Immunology, Oklahoma City, OK, USA; Heartland Veterinary Pathology Services, PLLC, Edmond OK, USA; Seattle Children’s Research Institute, Center for Integrative Brain Research, Seattle, WA, USA; Seattle Children’s Research Institute, Center for Immunity and Immunotherapies, Seattle, WA, USA; Departments of Pediatrics; Immunology, University of Washington, School of Medicine, Seattle, WA, USA; Benaroya Research Institute at Virginia Mason, Translational Research Program, Seattle, WA, USA

## Abstract

Type 1 diabetes (T1D) is an autoimmune disease in which pancreatic islet β-cells are attacked by the immune system, resulting in insulin deficiency and hyperglycemia. One of the top non-synonymous single-nucleotide polymorphisms (SNP) associated with T1D is in the interferon-induced helicase C domain-containing protein 1 (*IFIH1*), which encodes an anti-viral cytosolic RNA sensor. This SNP results in an alanine to threonine substitution at amino acid 946 (IFIH1^A946T^) and confers an increased risk for several autoimmune diseases, including T1D. We hypothesized that the *IFIH1^A946T^* risk variant, (*IFIH1^R^*) would promote T1D pathogenesis by stimulating type I interferon (IFN I) signaling leading to immune cell alterations. To test this, we developed *Ifih1^R^* knock-in mice on the non-obese diabetic (NOD) mouse background, a spontaneous T1D model. Our results revealed a modest increase in diabetes incidence and insulitis in *Ifih1^R^* compared to non-risk *Ifih1* (*Ifih1^NR^)* mice and a significant acceleration of diabetes onset in *Ifih1^R^* females. *Ifih1^R^* mice exhibited a significantly enhanced interferon stimulated gene (ISG) signature compared to *Ifih1^NR^*, indicative of increased IFN I signaling. *Ifih1^R^* mice exhibited an increased frequency of plasma cells as well as tissue-dependent changes in the frequency and activation of CD8^+^ T cells. Our results indicate that *IFIH1^R^* may contribute to T1D pathogenesis by altering the frequency and activation of immune cells. These findings advance our knowledge on the connection between the rs1990760 variant and T1D. Further, these data are the first to demonstrate effects of *Ifih1^R^* in NOD mice, which will be important to consider for the development of therapeutics for T1D.

## INTRODUCTION

Type 1 diabetes (T1D) is an autoimmune disease, where the pancreatic β-cells are attacked by the immune system, leading to insulin deficiency (1). Immune dysregulation is a hallmark of T1D. It is well known that cytotoxic CD8^+^ T cells implement β-cell destruction (2), but various other immune cells can drive β-cell targeted autoreactivity. Specifically, B cells have been shown to contribute to T1D pathogenesis by various mechanisms including antigen presentation, T cell co-stimulation, production of islet-reactive autoantibodies, and proinflammatory cytokines (3). Notably, children who develop T1D at a young age (<7 yo) show increased accumulation of CD20^+^ cells in islets and rapid β-cell loss compared to individuals developing T1D in their teen years (4, 5). These data suggest that B cells play a particularly pathogenic role in early onset T1D. Further, type I interferon (IFN I) has been shown to contribute to immune dysregulation. A surplus of IFN I has been associated with T1D and an IFN I signature is displayed in PBMCs from T1D patients prior to disease onset (6, 7). As IFN I is part of the anti-viral response, it is interesting that enterovirus infections, including coxsackievirus B (CVB), are associated with T1D (8), and CVB has been found to confer an increased risk for T1D (9). Importantly, the mechanisms linking viral infections and anti-viral immunity with diabetes and how these factors interact with genetic risk factors for diabetes remain unclear.

One gene that has been linked to T1D and is critical for anti-viral immunity is the cytosolic viral RNA sensor, interferon-induced helicase C domain-containing protein 1 (*IFIH1*), also known as MDA5 (10, 11). Upon recognition of dsRNA ligands, IFIH1 activates a signaling cascade leading to the transcription of pro-inflammatory cytokines, anti-viral genes, and IFN I (12, 13). A number of studies have demonstrated a link between IFIH1 and T1D pathogenesis (14-16). Decreased expression of IFIH1 was found to significantly decrease diabetes incidence (17) in NOD/ShiLtJ (NOD) mice, a spontaneous mouse model for T1D (18). NOD mice with an in-frame deletion in the *Ifih1* helicase domain exhibited decreased insulitis, IFN I production, and delayed onset of T1D (19). In contrast, the complete absence of *Ifih1* (*KO*) accelerated T1D in male mice, which was postulated to occur due to an inability of *Ifih1 KO* mice to produce sufficient myeloid-derived suppressor cells (19).

Further supporting the significance of IFIH1 in diabetes pathogenesis, a variant of *IFIH1*, rs1990760 was revealed as one of the top non-synonymous single-nucleotide polymorphisms (SNPs) associated with T1D (14). This variant results in an alanine to a threonine substitution at amino acid 946 (*IFIH1^A946T^*; *IFIH1*^R^), has been identified as a T1D susceptibility allele (14, 20), and is associated with an increased risk for multiple other autoimmune diseases. Consistent with the notion that elevated expression and/or activity of IFIH1 may drive diabetes progression, our previous work demonstrated the *Ifih1^A946T^* risk variant (*Ifih1^R^*) behaved as a gain-of-function variant, which elevated basal expression and poly(I:C)-induced levels of IFN I (21). In addition, *Ifih1^R^* accelerated streptozocin-induced diabetes incidence in mice expressing a diabetes risk variant of *Ptpn22* on a c57BL/6J background, supporting rs1990760 as a causal allele in T1D (21). Altogether, these findings highlight functional roles of IFIH1 in T1D pathogenesis. A recent study suggests that IFIH1 plays a role in immune cell compartments during T1D pathogenesis, particularly in the macrophages and T cells (19). However, how *IFIH1*^R^ may alter the immune system contributing to T1D pathogenesis remains unresolved. In this study, we aimed to decipher the mechanisms by which *Ifih1^R^* drives disease *in vivo*. To investigate the role of IFIH1^R^ in diabetes pathogenesis, the risk variant rs1990760 was generated on the NOD background. Findings from the present study reveal a presence of an ISG signature and immune cell activation in NOD mice expressing *Ifih1^R^*. Further, an acceleration in disease in female NOD mice expressing *Ifih1^R^* was also observed. Collectively, our findings identify potential mechanisms by which *Ifih1^R^* contributes to diabetes pathogenesis *in vivo* and demonstrate how the *Ifih1^R^* NOD model provides an opportunity to investigate the connection between the human risk variant, rs1990760 and T1D.

## MATERIAL AND METHODS

### Ethics statement

Procedures and experiments performed on all mice followed the Institutional Animal Care & Use Committee guidelines of the Oklahoma Medical Research Foundation and Seattle Children’s Research Institute.

### Animals

To generate the *Ifih1*^R^ allele on the NOD background, a NOD embryonic stem (ES) cell line was first derived from the inner cell mass (ICM) of naturally mated NOD/ShiLtJ (Jackson Laboratory, Bar Harbor, ME, Stock No 001976) animals. Blastocysts were plated onto mitomycin C-inactivated CD1 (Charles River Laboratories, Hollister, CA) mouse embryonic fibroblasts (MEFs). After the blastocysts hatched, outgrowths were picked, lightly mechanically disaggregated and transferred to feederless collagen-coated plates (22), but maintained in 3i culture medium (23). A plasmid construct designed to generate a A946T mutation in exon 15 of *Ifih1* by homologous recombination and *lox*P sites for potential lineage-specific deletion in NOD mice was generated by Biocytogen (21). The *lox*P sites were not used for lineage-specific deletion in this study. One sub-confluent 12-well dish of low passage feederless NOD embryonic stem (ES) cells was transfected with 250ng of each of 2 pX330 constructs (containing Cas9 and 1 sgRNA) and 500 ng of the plasmid template with FuGene HD transfection reagent (Promega Corporation, Madison, WI) in the recommended conditions. After 24h, the cells were replated with medium containing 800µg/mL G418. Surviving colonies were picked on days 7–8. Individual colonies were dissociated with trypsin and re-seeded onto collagen-coated 48-well plates containing 3i media with G418. After a clone became confluent, it was trypsinized and 10% of the cells were replated onto a 24 well plate. The remaining 90% of cells were submitted for gDNA screening by PCR to confirm correct integration. Approximately 10-14 cells from successfully targeted clones were injected into c57BL/6J blastocysts, which were then transferred into the uteri of pseudopregnant CD-1 females. Microinjections were performed under an inverted microscope (Olympus IX71) equipped with Eppendorf micromanipulators. The procedure was carried out at room temperature in DME/High Glucose medium (HyClone Laboratories, Logan, UT). After microinjection, embryos were immediately transferred into pseudopregnant CD-1 females or cultured in BMOC medium (Thermo Fisher Scientific, Waltham, MA) at 37.5°C in 5% CO2 in air overnight and transferred the next day. Three male chimeric founder mice (F0) were recovered and bred to NOD females, of which one exhibited transmission to the next generation (F1) based on genotyping. Animals were housed and maintained under a 14:10-h (light-dark) cycle. Mice heterozygous for the *Ifih1* risk allele (*Ifih1^NR/R^*) were transferred to the Oklahoma Medical Foundation (OMRF) and bred together to establish the mouse colony at OMRF. The sequence of *Ifih1^R^* and *Ifih1^NR^* in the NOD mice were confirmed using purified tail DNA from 4 different mice (2 *Ifih1^R^* and 2 *Ifih1^NR^* mice) and amplifying an 816 base pair region of the *Ifih1* gene using the A946 F1 and A946 R1 primers (Supplementary Table 1). The PCR product was purified using the Zymo Research Genomic DNA Clean & Concentrator kit and sequenced using the A946 R2 primer (Supplementary Table 1). Sequence analysis was performed using DNASTAR Lasergene17 software.

### Diabetes Incidence

Blood glucose was measured from the lateral saphenous veins of mice every two weeks using the Contour NEXT EX Blood Glucose Monitoring System (Ascencia) from 10 to 32 weeks of age. Mice with two consecutive readings of ≥ 250mg/dL were considered diabetic. Mice were euthanized if they had a reading ≥500mg/dL.

### Flow Cytometry

The spleen and non-draining lymph nodes (mesenteric, axillary, and inguinal) were harvested from NOD *Ifih1^R/R^* and *Ifih1^NR/NR^*mice. Spleens and non-draining lymph nodes were digested in 3 mL of 1.6 mg/mL collagenase Type IV (Worthington Biochemical), 0.1 mg/mL DNase I (Thomas Scientific) and complete cell culture medium (RPMI supplemented with 10% fetal bovine serum (FBS), sodium pyruvate, Glutamax, non-essential amino acids, and 2-mercaptoethanol) at 37°C in a Jitterbug Microplate Shaker (Boekel Scientific). After 30 minutes (min, for the spleen) and 15 min (for the pooled lymph nodes), cell dissociation buffer (Gibco) was added, and tissues were incubated for an additional 5 min. Tissues were mashed through a 40μm nylon cell strainer to obtain a single-cell suspension. Splenocytes were treated with lysing Buffer (BD Biosciences) for 2 min, and then washed twice with cell culture media. Lymph node cells were washed once. Pancreas tissue was minced with a blade and digested in 1mg/mL of collagenase P (Millipore Sigma, Rockville, MD) and 0.1mg/mL of DNAse (Millipore Sigma) for 40 min with shaking at 37°C. Following digestion, tissues were placed on ice for 10 min and then filtered through 100μm nylon mesh. Following, tissues were washed with FACS buffer (DMEM with 5% FBS and 10mM HEPES, Gibco) treated with 1mL of lysing buffer for 3 min and washed again with FACS buffer. Pancreatic draining lymph nodes were directly mashed through a 40μm nylon cell strainer to obtain a single-cell suspension and washed once with cell culture media. Single cell suspensions were counted using Precision Count Beads (Biolegend) and stained with 20 μg/ml DAPI to determine cell viability. Spleen and non-draining lymph node single cell suspensions were seeded at 2×10^6^ cells/well and 1-2×10^6^ cells/well, respectively into 96-well plates. For pancreatic lymph nodes, the entire single cell suspension obtained from each mouse was seeded into one well of a 96-well plate. For pancreas samples, 1mL (out of a 5mL suspension) was transferred to a FACS tube for staining. Cells were washed with phosphate-buffered saline (PBS) and incubated with FcR blocking reagent (Miltenyi Biotec) prior to incubating with antibodies (Supplementary Table 2). After staining with antibodies, cells were washed twice with PBS. Prior to staining samples for detection of intracellular cytokines, cells were incubated for 5 hr with and without 50ng/mL phorbol 12-myristate 13 acetate (PMA) and 1μg/mL ionomycin to stimulate cytokine production in T cells as well as GolgiPlug (BD Biosciences) to retain cytokines in the cytoplasm. After staining extracellular antigens, cells were then fixed and permeabilized using the Cytofix/Cytoperm kit (BD Biosciences) following manufacturer recommendations. Staining for intracellular cytokines was performed on the fixed/permeabilized cells. All samples were run on an Aurora Flow Cytometer (Cytek) and data was analyzed using FlowJo Prism software. Near IR fluorescent reactive dye was used to exclude dead cells in the analyses. As a quality control measure, pancreas samples with < 2% viable cells were excluded from analysis.

### qRT-PCR

RNA from 5×10^6 splenocytes was extracted with the RNeasy Mini Kit (BioRad) and converted to cDNA using the Maxima First Strand cDNA Synthesis Kit (ThermoFisher Scientific). Quantitative reverse transcriptase polymerase chain reation (RT-qPCR) using iTaq Universal Syber Green Supermix (BioRad) and respective primers (Supplementary Table 1) was performed on cDNA to quantify the expression of *Ifih1, Mx1, Ifit1, Hprt, and Oas1* genes. The plates were run on a Roche LightCycler 480 II. For Δ*C*_t_ values, gene expression was normalized to *Hprt*. Fold differences in mRNA expression (relative expression) were determined using the equation [2^(-ΔΔCt^)].

### Histology

Pancreases were fixed in 10% neutral buffered formalin for 24 hours (hr) and submitted to the OMRF Imaging Core Facility for paraffin processing, sectioning (on long axis) and hematoxylin and eosin (H&E) staining. Representative images were captured on a Zeiss Axioscan7. Insulitis was scored by an observer blinded to experimental details. For each mouse, 10 non-overlapping fields from 3 sections 100 microns apart were scored at 20X magnification. A grading score of 0-4 was used: 0=no infiltration, 1=per-insulitis, 2=<25% islet infiltration, 3=25-75% islet infiltration, and 4=>75% islet infiltration. Scores were excluded if less than 10 islets/section could be identified. The number of islets in each of 10 randoms fields were counted from 1 section per mouse. The average insulitis score, number of islets, and insulitis/islet (assessed from 1 section/mouse) per slide was plotted.

### Immunofluorescent staining

The OMRF Imaging Core Facility performed Immunofluorescent (IF) staining on formalin-fixed paraffin embedded pancreas tissues (see Histology section). Pancreas sections were stained with rat anti-mouse B220/CD45R (catalog # 14-0452-82, ThermoFisher Scientic) together with rabbit anti-insulin (catalog # 3014, Cell Signaling Technology) primary antibodies. The secondary antibodies used were AlexaFluor 488 donkey anti-rabbit (Jackson ImmunoResearch, West Grove, PA) and AlexaFluor 647 donkey anti-rat (Jackson ImmunoResearch). For each mouse, all islets from one section (11-58mm^2^, mean 32mm^2^) were analyzed using Image J software. The insulin-positive regions of the islets and whole islets were each outlined to measure the respective areas and calculate the percentage of insulin covered area per islet. To determine the percentage of B cell covered area per islet, the threshold tool in Image J was used to eliminate background signal and measure the percent area covered by B cells for each islet (whole islets were outlined using the region of interest manager in Image J). Mean insulin fluorescence intensity was also determined using the same process with the threshold and region of interest tools of Image J. Representative images were captured on a Zeiss Axioscan7 at 40X magnification. The average scores/islet for each mouse were plotted.

### Statistical analysis

Statistical analysis was performed using GraphPad Prism 9.0. All data are expressed as mean ± SEM with p values of <0.05 considered significantly different. The Gehan-Breslow-Wilcoxon analysis was used for assessing differences in Kaplan-meier curves for diabetes incidence. Differences between *Ifih1*^R/R^ and *Ifih1*^NR/NR^ mice were determined using student’s unpaired *t*-tests.

## RESULTS

### Generation of *NOD.Ifih1^R^* mice

To determine the role of the *IFIH1* risk variant in the immune cell compartments during diabetes pathogenesis *in vivo*, we generated a mouse strain with the *Ifih1* risk allele that precisely mimicked the haplotype of two human T1D risk variants, rs3747517 (*IFIH1^H843R^*) and rs1990760 (*IFIH1^A946T^*), that were previously shown to be significantly associated with T1D (21). NOD mice naturally contain an arginine at amino acid 843 in the *Ifih1* gene (risk allele; *Ifih1^R843^*), allowing us to study the contribution of *Ifih1^A946T^* to the risk haplotype on the NOD background. We used CRISPR/Cas9 editing to establish mice expressing the *Ifih1^A946T^* risk allele on the NOD/ShiLtJ (NOD) background strain resulting in the risk haplotype encoded on the NOD background consisting of an arginine at position 843 and threonine at position 946 of *Ifih1* mouse gene (called “*NOD.Ifih1^R843-T946”^* (or simply *NOD.Ifih1^R^,* here)) (**Fig. 1A-B).** Mice homozygous for the risk *Ifih1* allele (*NOD.Ifih1^R/R^)* did not exhibit significant differences in *Ifih1* mRNA levels at 14 weeks of age **(Fig S1A-B)**. Further, *NOD.Ifih1^R/R^* mice were viable and fertile **(Fig. 1C)**. No differences were detected in body weights between homozygous non-risk *Ifih1* (homozygous genotype of *NOD.Ifih1^R843-A946^* (or simply *NOD.Ifih1^NR/NR^,* here)) and *NOD.Ifih1^R/R^* mice at 14 weeks of age **(Fig. S2A-B)**.

**Figure 1.**
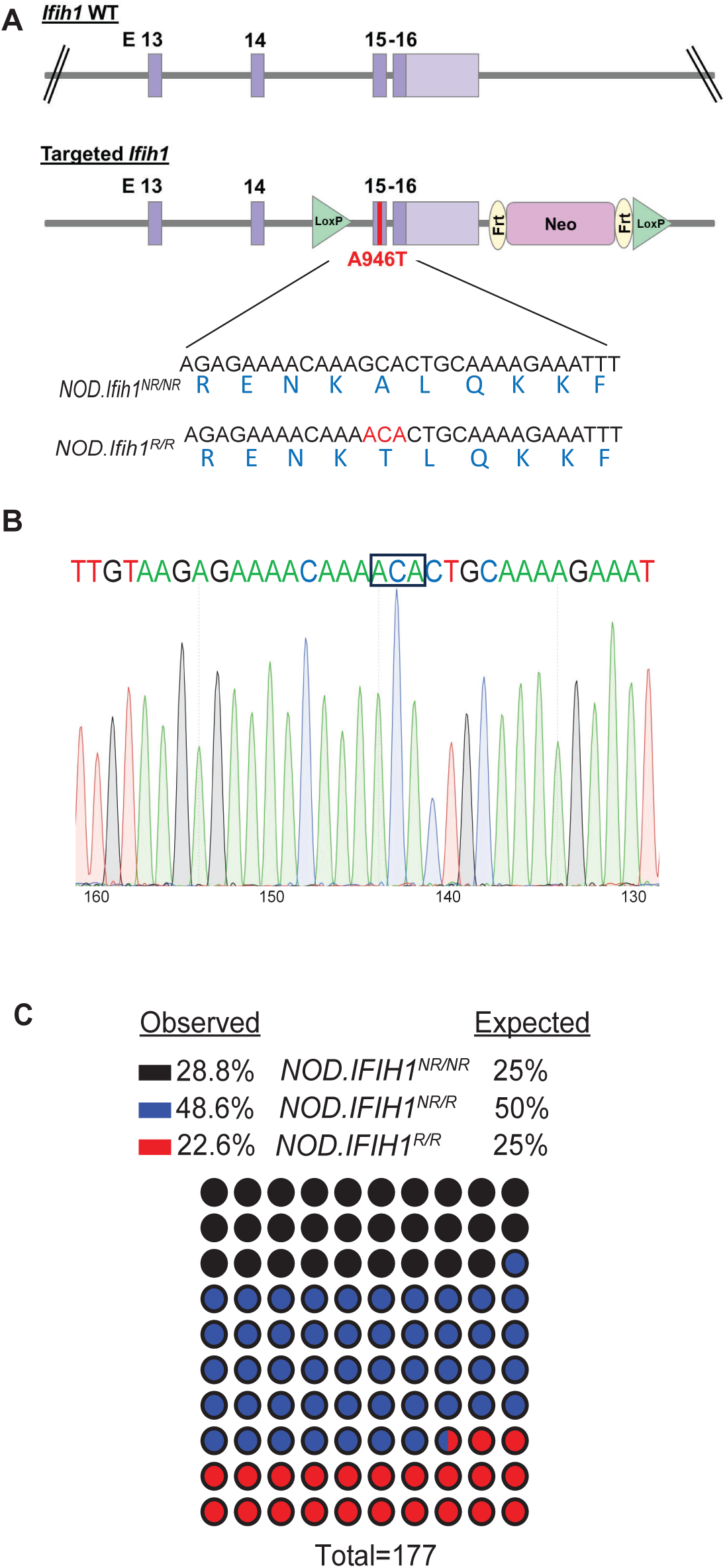
Generation of *NOD.Ifih1^R^* mice. **(A)** Illustration of the location of the point mutation in *Ifih1* at exon 15 as well as *lox*P sites using the CRISPR/Cas9 system. The point mutation is indicated in red **(B)** Sequence of the *Ifih1* mutation. The point mutation is boxed. **(C)** Genotypes of mice (*n*=177) derived from the interbreeding of NOD.*Ifih1^NR/R^* heterozygous mice.

### *NOD.Ifih1^R/R^* mice exhibit an elevated ISG signature

We have previously shown that c57BL/6J mice expressing *Ifih1^R^*exhibit an elevated basal ISG signature (21). Due to our previous identification of *Ifih1^R^* as a gain-of-function variant (21), we hypothesized that *NOD.Ifih1^R/R^*mice would also display an elevated ISG signature. Quantitative reverse transcriptase polymerase chain reaction (qRT-PCR) revealed a significant increase in *Mx1, Ifit1,* and *Oas1* in the spleen of *NOD.Ifih1^R/R^* compared to *NOD.Ifih1^NR/NR^* mice. Importantly, these ISGs were significantly upregulated in both male and female *NOD.Ifih1^R/R^* mice at 14 weeks of age **(Fig. 2A-B)**. Thus *NOD.Ifih1^R/R^* mice exhibited a similar increased basal ISG signature as in c57BL/6J *Ifih1^R/R^* mice.

**Figure 2.**
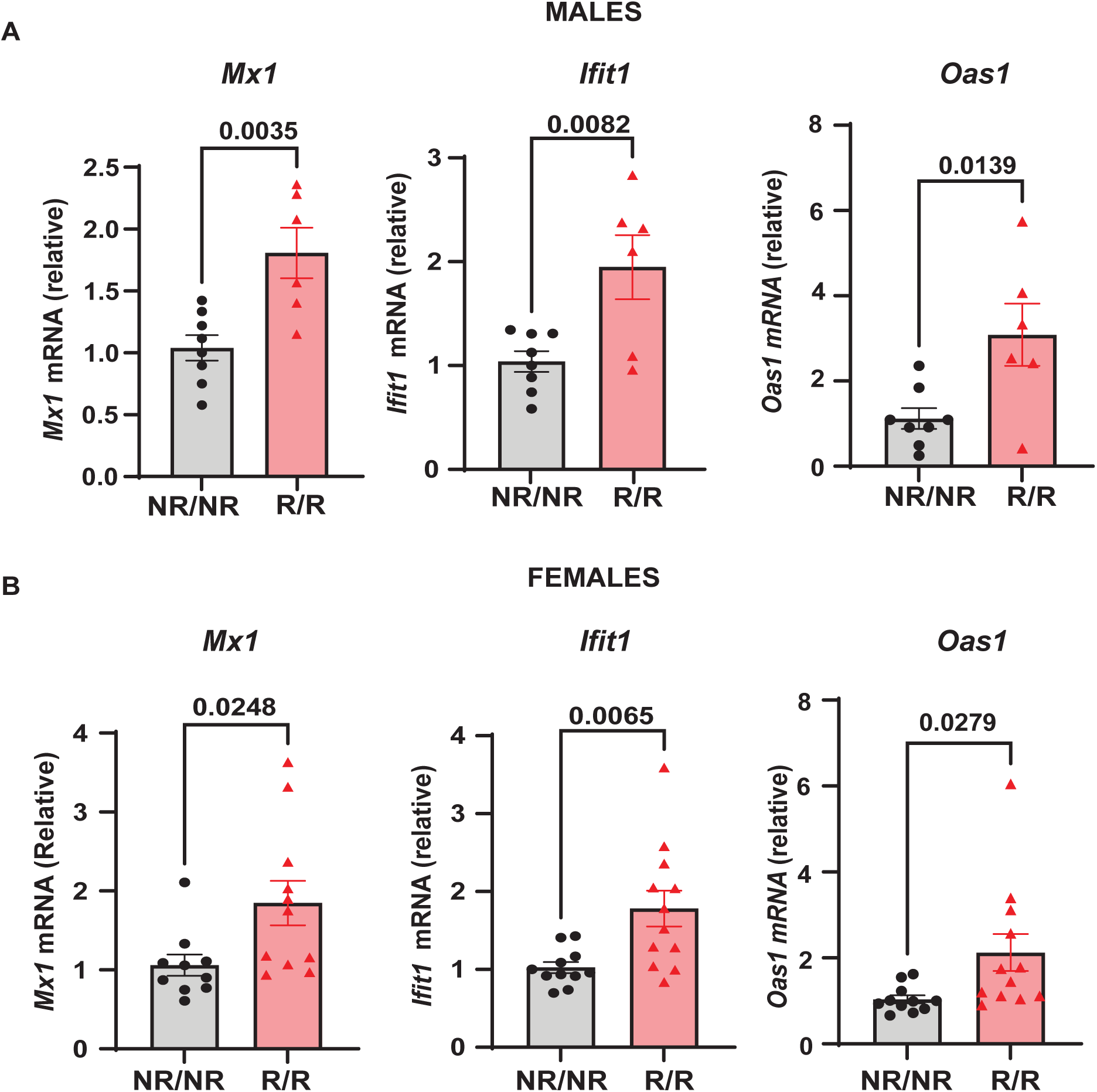
*NOD.Ifih1^R^* mice exhibit an enhanced ISG signature. The relative mRNA expression of ISGs, including *Mx1*, *Ifit1*, and O*as1* in 14-week-old **(A)** males (*n*=8 *NOD.Ifih1^NR/NR^*, *n*=6 *NOD.Ifih1^R/R^*) and **(B)** females (*n*=11 *NOD.Ifih1^NR/NR^*, *n*=12 *NOD.Ifih1^R/R^*) separated. P values were determined using student’s unpaired *t* tests. All data are mean ± SEM.

### *NOD.Ifih1*^R^ alters the frequency and activation of CD8^+^ T cells in a tissue-dependent manner

Altered activation and composition of immune cells play a major role in pancreatic β-cell loss. We proposed that *NOD*.*Ifih1^R/R^* would change immune cell populations through the elevated IFN signature exhibited in *NOD.Ifih1^R/R^*mice. Due to the prominent role of autoreactive T cells in T1D pathogenesis, we first assessed spleens and lymph nodes from *NOD.Ifih1^R/R^* mice for changes in T cell frequency and activation. No differences were observed in the spleen weights (**Fig. S3A**), total numbers of cells, (**Fig. S3B-C**) or cell viability in spleens or non-draining lymph nodes (**Fig S3D-E**). In addition, *NOD*.*Ifih1^R/R^* mice showed no difference in the frequency or numbers of CD4^+^ T cells **(Fig. S4A-D)**. Interestingly, we detected a significant increase in the frequency and total number of splenic CD8^+^ T cells in *NOD*.*Ifih1^R/R^*mice **(Fig. 3A-B).** However, the percentage of splenic CD8^+^ T cells that were CD69^+^ was decreased **(Fig. 3C-D)**. In contrast to the spleens, we did not observe an increase in the frequency or total number of CD8^+^ T cells in *NOD*.*Ifih1^R/R^* non-draining lymph nodes **(Fig. 4A-B).** However, we detected an increased frequency of CD69^+^CD8^+^ T cells in *NOD*.*Ifih1^R/R^* females alone **(Fig. 4C-E)**. To further evaluate CD8^+^ T cell activation, we assessed CD8^+^ T cells for intracellular production of IFN-γ, a major pro-inflammatory cytokine produced by activated CD8^+^ T cells (24). Cells were treated with phorbol 12-myristate 13 acetate (PMA) and ionomycin, which stimulate IFN-γ production in T cells. Our results showed an increase in the frequency of IFNγ^+^CD8^+^ central memory and effector memory T cells in non-draining lymph nodes of *NOD*.*Ifih1^R/R^*compared to *NOD.Ifih1^NR/NR^* mice **(Fig. 5A-C)**. No difference was observed in the frequency of splenic IFNγ^+^CD8^+^ T cells **(Fig. S5A-C)**. Therefore, *Ifih1^R^* exerted differential effects on CD8^+^ T cell activation in the spleen compared to the lymph nodes. Altogether, these findings suggest that *NOD*.*Ifih1^R/R^* may impact diabetes pathogenesis by altering CD8^+^ T cell-dependent immune responses in a tissue-dependent manner.

**Figure 3.**
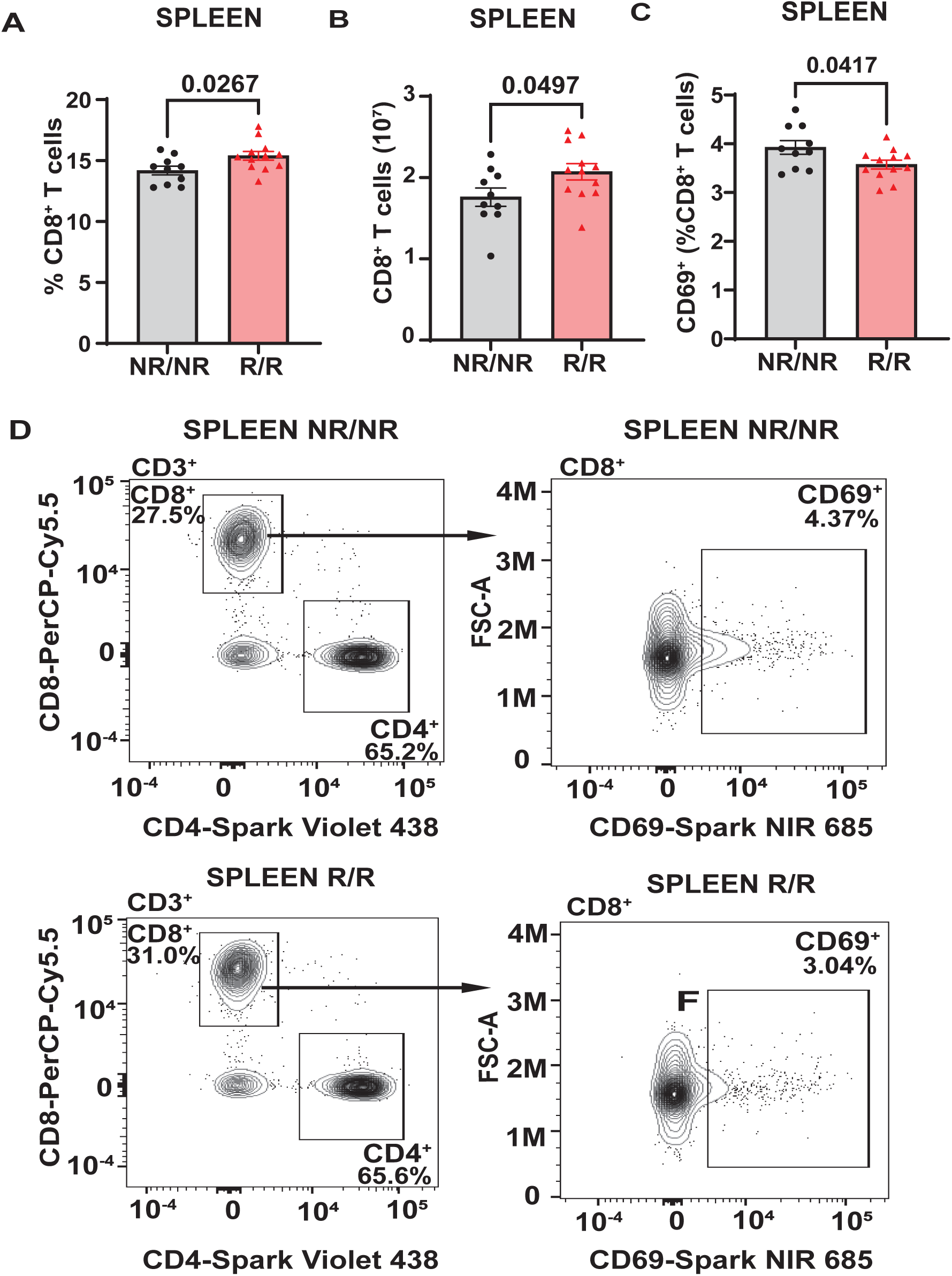
*NOD.Ifih1^R^* increases the frequency of splenic CD69^-^ CD8^+^ T cells. **(A-B)** CD8^+^ T cells in the spleen as **(A)** frequency of live cells and **(B)** total CD8^+^ T cell numbers in *NOD.Ifih1^NR/NR^*compared to *NOD.Ifih1^R/R^* mice (*n*=10 *NOD.Ifih1^NR/NR^*, *n*=12 *NOD.Ifih1^R/R^*). **(C)** The splenic frequency of CD69^+^CD8^+^ T cells (CD3^+^CD8^+^CD69^+^) in 14-week-old *NOD.Ifih1^NR/NR^* compared to *NOD.Ifih1^R/R^* mice (*n*=10 *NOD.Ifih1^NR/NR^*, *n*=12 *NOD.Ifih1^R/R^*). **(D)** Representative gating of CD69^+^ CD8^+^ T cells (CD3^+^CD8^+^CD69^+^) from 14-week-old *NOD.Ifih1^NR/NR^* and *NOD.Ifih1^R/R^* mice by flow cytometry. P values were determined using student’s unpaired *t* tests. All data are mean ± SEM.

**Figure 4.**
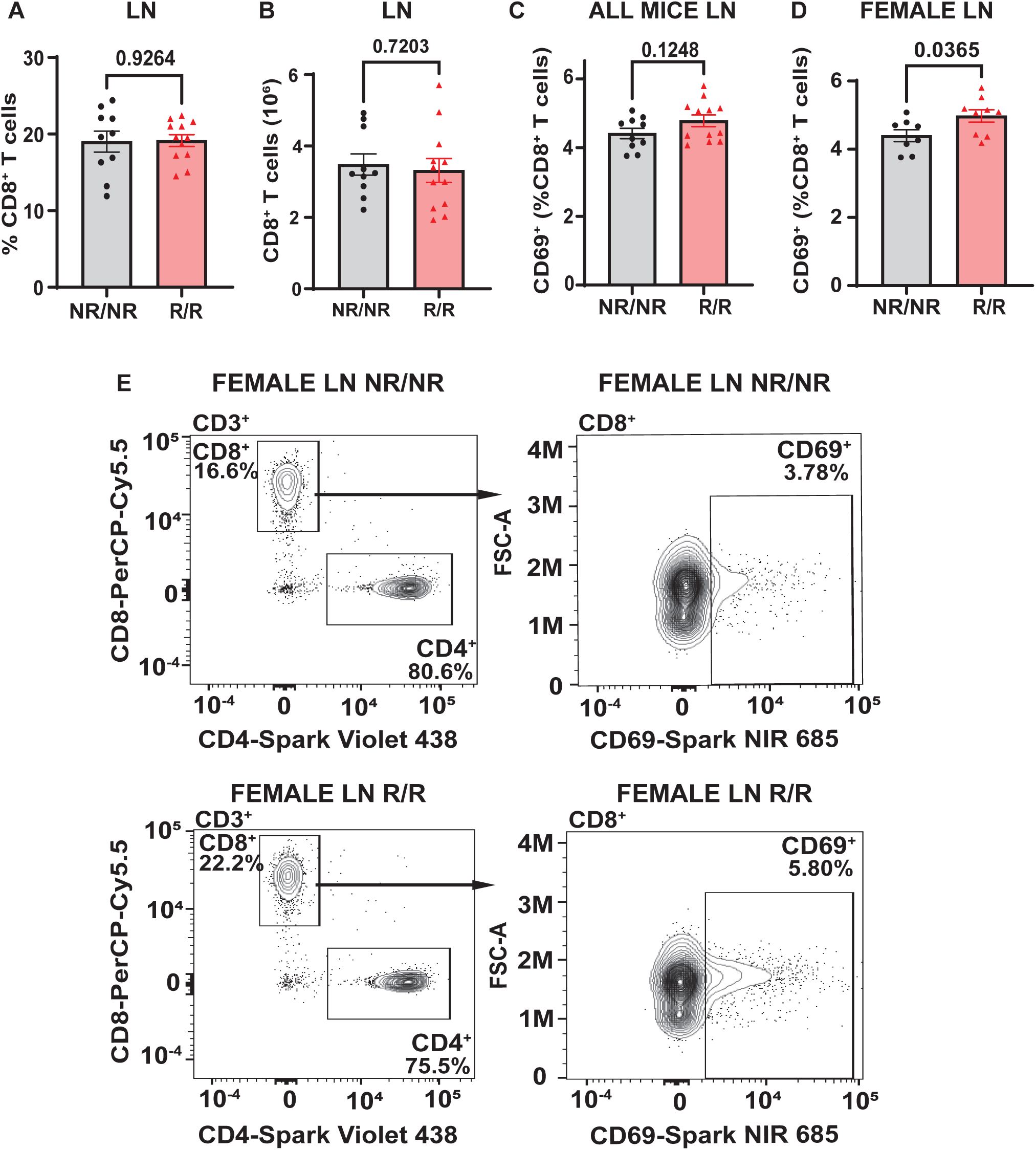
*NOD.Ifih1^R^* increases the frequency of CD69 on CD8^+^ T cells in female non-draining lymph nodes. **(A-B)** CD8^+^ T cells in the lymph nodes (LN) as **(A)** frequency of live cells and **(B)** total CD8^+^ T cell numbers in *NOD.Ifih1^NR/NR^* compared to *NOD.Ifih1^R/R^* mice (*n*=10 *NOD.Ifih1^NR/NR^*, *n*=12 *NOD.Ifih1^R/R^*). **(C-D)** The frequency of CD69^+^ CD8^+^ T cells (CD3^+^CD8^+^CD69^+^) from lymph nodes of **(C)** all 14-week-old *NOD.Ifih1^NR/NR^* compared to *NOD.Ifih1^R/R^*mice (*n*=10 *NOD.Ifih1^NR/NR^*, *n*=12 *NOD.Ifih1^R/R^*) and **(D)** female *NOD.Ifih1^NR/NR^* compared to *NOD.Ifih1^R/R^* mice (*n*=8 *NOD.Ifih1^NR/NR^*, *n*=9 *NOD.Ifih1^R/R^*). **(E)** Representative gating of CD69^+^ CD8^+^ T cells (CD3^+^CD8^+^CD69^+^) from lymph nodes of 14-week-old female *NOD.Ifih1^NR/NR^* and *NOD.Ifih1^R/R^* mice by flow cytometry.

**Figure 5.**
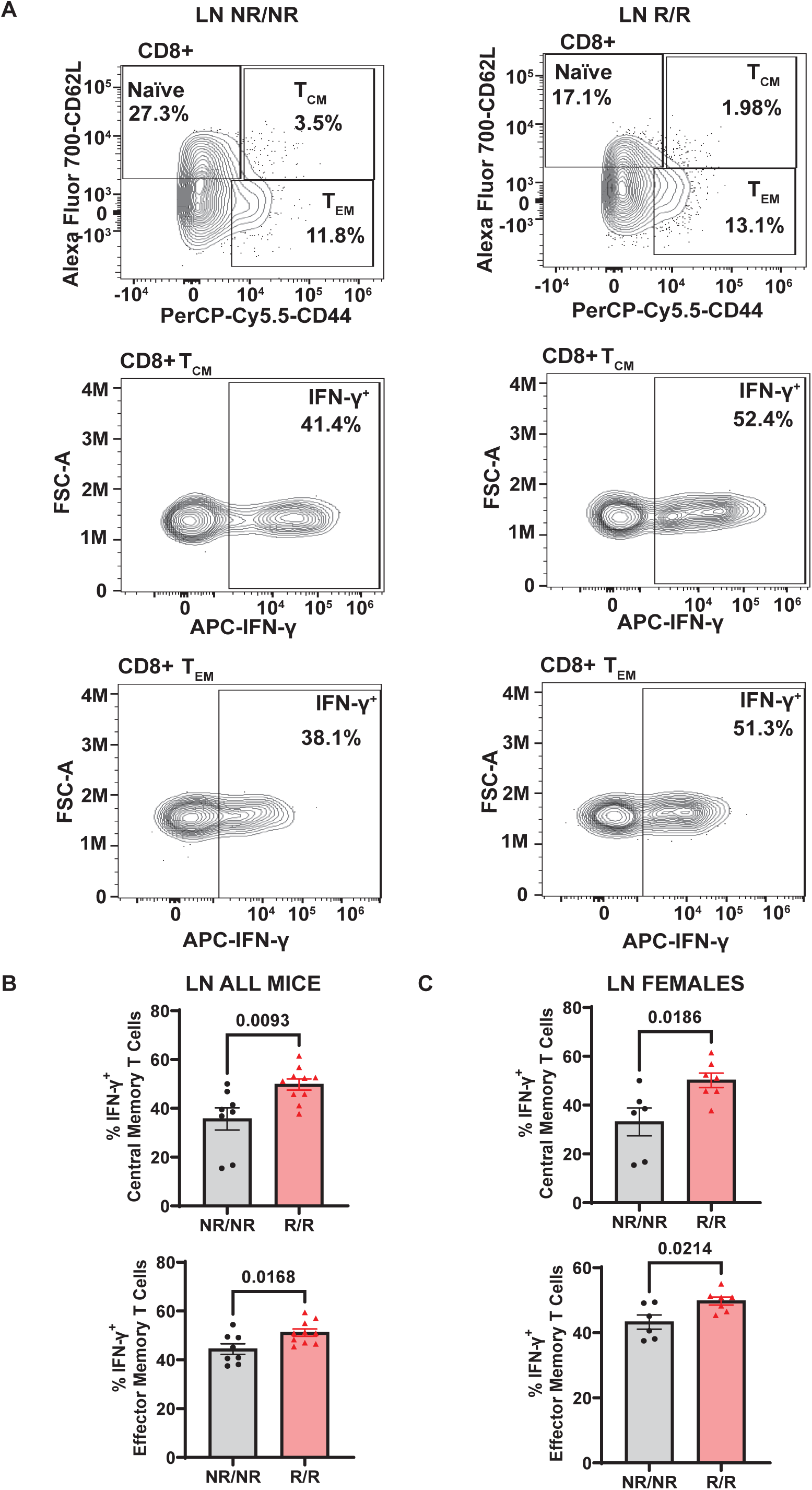
*NOD.Ifih1^R^* increases the frequency of IFN-γ^+^CD8^+^ central memory and effector memory T cells in non-draining lymph nodes. **(A)** Representative gating of CD8^+^ Naïve (CD62L^+^), central memory (CD62L^+^CD44^+^, TCM), effector memory (CD62L^-^CD44^+^, TEM), and IFN-γ^+^CD8^+^ central memory and effector memory T cells from PMA/ionomycin-treated non-draining lymph nodes. **(B-C)** The frequency of IFN-γ^+^ CD8^+^ central memory and effector memory T cells from PMA/ionomycin-treated non-draining lymph nodes from **(B)** all mice (*n*=8 *NOD.Ifih1^NR/NR^*, *n*=10 *NOD.Ifih1^R/R^*) and **(C)** females only (*n*=6 *NOD.Ifih1^NR/NR^*, *n*=7 *NOD.Ifih1^R/R^)*. P values were determined using student’s unpaired *t* tests. All data are mean ± SEM.

### *NOD.Ifih1*^R^ increases B cell activation in lymph nodes

In addition to T cells, we focused on B cells due to their critical role in T1D pathogenesis in both humans and mice (5, 25). Since IFN I are known to drive B cell activation in a multipotent manner (26) and *NOD.Ifih1^R/^*^R^ exhibited an elevated ISG signature, we assessed for changes in B cells. By flow cytometry, there were no differences in the frequency or total number of B cells in spleens **(Fig. S6A-B)** or non-draining lymph nodes **(Fig. S6C-D)**. However, the frequency of plasma cells was significantly increased in the non-draining lymph nodes of *NOD.Ifih1^R/R^*compared to *NOD.Ifih1^NR/NR^* mice **(Fig 6A-B)**. In addition, non-draining lymph nodes from *NOD.Ifih1^R/R^* mice showed a trend for an increased frequency of CD80^+^ B cells **(Fig. 6C-D)**. As CD80 is important for B-T cell interactions and CD80 deficiency has been found to decrease the frequency of long-lived plasma cells (27), the increased frequency of CD80^+^ B cells in *NOD.Ifih1^R/R^* mice may have facilitated increased B-T cell interactions and plasma cell formation or survival. To further assess B cell activation, we examined levels of an anti-inflammatory cytokine, IL-10, in B cells and found a significantly decreased frequency of IL-10^+^ cells that were positive for the plasma cell marker, CD138 in spleens from *NOD.Ifih1^R/R^* compared to *NOD.Ifih1^NR/NR^* mice **(Fig. 6E-F)**. A similar trend for decreased IL-10 frequency was observed in CD138^+^ cells from non-draining lymph nodes of *NOD.Ifih1^R/R^* compared to *NOD.Ifih1^NR/NR^* mice **(Fig. 6G-H)**. As previous studies have indicated that IL-10 secreting plasma cells or plasmablasts may play a regulatory role in autoimmune diseases (28, 29), a decrease in IL-10 production by *NOD.Ifih1^R/R^* CD138^+^ cells may support enhanced inflammation during diabetes. Despite the increase in plasma cell frequency observed in *NOD.Ifih1^R/R^*mice, no differences in insulin autoantibody production were detected between *NOD.Ifih1^R/R^* and *NOD.Ifih1^NR/NR^* male or female mice at age 14 weeks of age **(Fig. S7A)**. Additionally, no differences were revealed in the levels of Smith ribonucleoprotein (smRNP) or dsDNA autoantibodies (**Fig. S7B-E**), which were found to be elevated by *Ifih1^R^* in c57BL/6J mice given adoptive transfer of BM12 CD4^+^ T cells in our previous study (21). These findings support the notion that *Ifih1^R^* increases B cell activation in females and promotes disease in female *NOD.Ifih1^R/R^* mice by methods other than autoantibodies.

**Figure 6.**
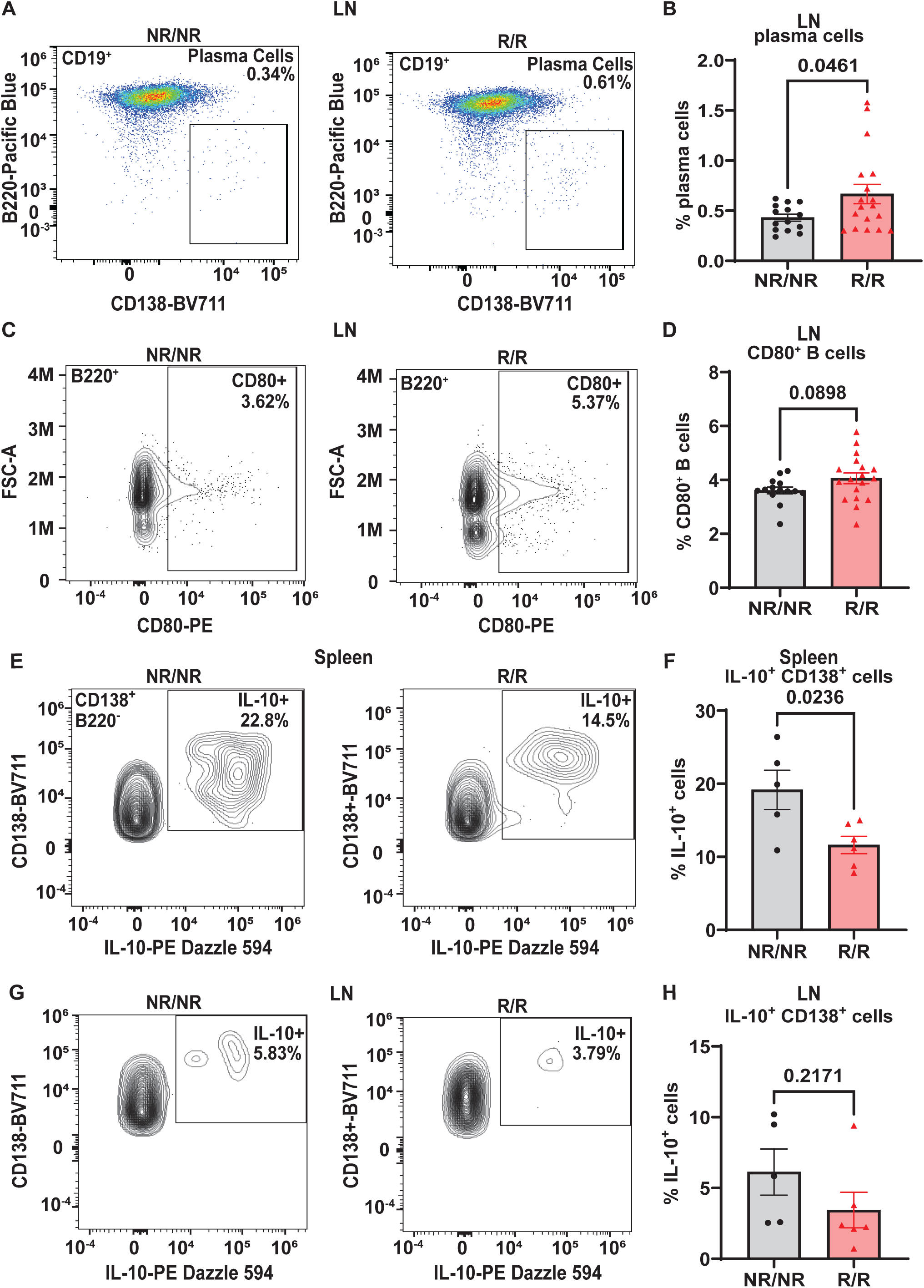
*NOD.Ifih1^R^* increases B cell activation in non-draining lymph nodes and decreases the frequency of splenic IL-10^+^CD138^+^ cells. (A) Representative gating of plasma cells (CD19^+^B220^-^CD138^+^) from lymph nodes of 14-week-old *NOD.Ifih1^NR/NR^* and *NOD.Ifih1^R/R^* mice by flow cytometry. **(B)** The frequency of plasma cells in lymph nodes of 14-week-old mice (*n*=14 *NOD.Ifih1^NR/NR^*, *n*=18 *NOD.Ifih1^R/R^*). **(C)** Representative gating of CD80^+^ B cells (B220^+^CD80^+^) from lymph nodes of 14-week-old *NOD.Ifih1^NR/NR^* and *NOD.Ifih1^R/R^* mice by flow cytometry. **(D)** The frequency of CD80^+^ B cells in lymph nodes of 14-week-old mice (*n*=14 *NOD.Ifih1^NR/NR^*, *n*=18 *NOD.Ifih1^R/R^*). **(E)** Representative gating of IL-10^+^ B220^-^CD138^+^ cells from spleens of 14-week-old *NOD.Ifih1^NR/NR^* and *NOD.Ifih1^R/R^* mice by flow cytometry. **(F)** The frequency of IL-10^+^ B220^-^CD138^+^ cells from spleens of 14-week-old mice (*n*=5 *NOD.Ifih1^NR/NR^*, *n*=6 *NOD.Ifih1^R/R^*). **(G)** Representative gating of IL-10^+^ B220^-^ CD138^+^ from lymph nodes of 14-week-old *NOD.Ifih1^NR/NR^* and *NOD.Ifih1^R/R^* mice by flow cytometry. **(H)** The frequency of IL-10^+^ B220^-^CD138^+^ cells from lymph nodes of 14-week-old mice (*n*=5 *NOD.Ifih1^NR/NR^*, *n*=6 *NOD.Ifih1^R/R^*). P values were determined using student’s unpaired *t* tests. All data are mean ± SEM.

### *NOD.Ifih1*^R^ increases diabetes incidence and insulitis

Due to the differences in the immune system, we next wanted to determine the effects of *Ifih1*^R^ on diabetes incidence. Blood glucose levels were assessed every two weeks between 10-32 weeks of age in mice expressing the *NOD.Ifih1^NR/NR^* or *NOD.Ifih1^R/R^.* Mice were considered diabetic if they showed 2 consecutive blood glucose readings of ≥250 mg/dL. The rate of diabetes was modestly increased in *NOD.Ifih1^R/R^* compared to *NOD.Ifih1^NR/NR^*mice **(Fig. 7A)**. The incidence of diabetes was significantly accelerated in female *NOD.Ifih1^R/R^* compared to *NOD.Ifih1^NR/NR^*mice **(Fig. 7B)**. Strikingly, by 16 weeks of age, greater than 70% of *NOD.Ifih1^R/R^* female mice became diabetic, whereas less than 25% of *NOD.Ifih1^NR/NR^* females were diabetic at this age. In contrast, male *NOD.Ifih1^R/R^* mice did not exhibit a significant acceleration in diabetes incidence. *NOD.Ifih1^R/R^* males showed a modest, yet insignificant increase in diabetes incidence by age 30 weeks **(Fig. 7C)**. Altogether, these findings indicate a sex-dependent effect of *NOD.Ifih1^R^* in driving diabetes pathogenesis.

**Figure 7.**
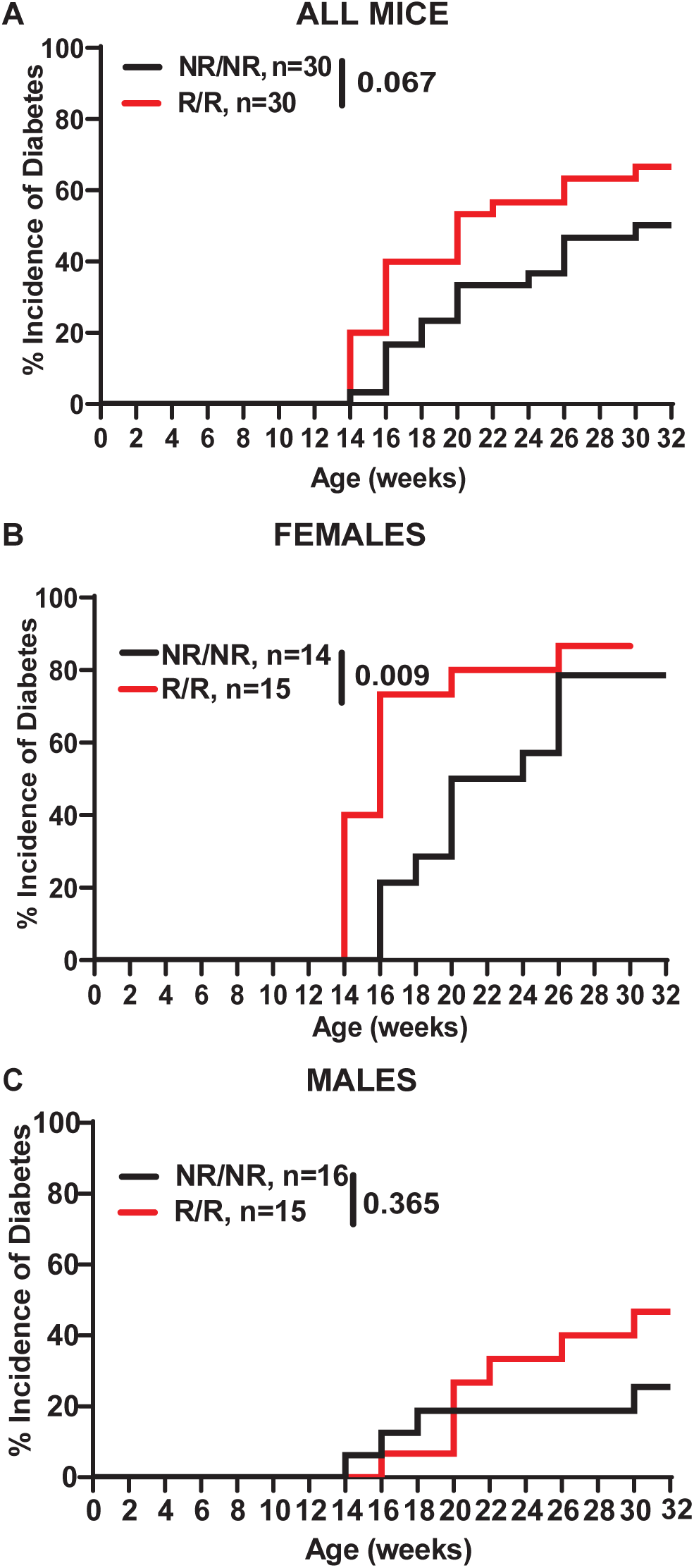
*NOD.Ifih1^R^* accelerates diabetes incidence in females. **(A-C)** Diabetes incidence in **(A)** total mice *(n*=30 per genotype), **(B)** females only (*n*=14 *NOD.Ifih1^NR/NR^*, *n*=15 *NOD.Ifih1^R/R^*), and **(C)** males only (*n*=16 *NOD.Ifih1^NR/NR^*, *n*=15 *NOD.Ifih1^R/R^*), and displayed using Kaplan Meier survival curves. P values for diabetes incidence in A-C were determined by performing Gehan-Breslow-Wilcoxon tests.

Prior to the onset of hyperglycemia and overt diabetes, immune cells infiltrate pancreatic islets and attack islet β-cells. In NOD mice, insulitis begins to occur as early as 3-4 weeks of age and continues to worsen until clinical symptoms of diabetes manifest, which generally occurs between 14-25 weeks of age (30). To determine the effect of *Ifih1^R^*on the magnitude of insulitis, we assessed insulitis in *NOD.Ifih1^R/R^*and *NOD.Ifih1^NR/NR^* mice at age 14 weeks, an early timepoint in disease pathogenesis. Our results revealed significantly elevated insulitis in *NOD.Ifih1^R/R^* compared to *NOD.Ifih1^NR/NR^* mice **(Fig. 8A-F)**. Interestingly, although acceleration of diabetes occurred only in female *NOD.Ifih1^R/R^*mice, only male *NOD.Ifih1^R/R^* exhibited a statistical increase in inflammation at this time point as compared to *NOD.Ifih1^NR/NR^*mice **(Fig. 8C-F)**. Further, no differences were observed in the numbers of pancreatic islets in *NOD.Ifih1^R/R^* compared to *NOD.Ifih1^NR/NR^* mice at this time point **(Fig. 8G-H).** Overall, these results demonstrate that while *Ifih1^R^* accelerates diabetes onset in female mice, this accelerated process is associated with only modest changes in pancreatic immune cell infiltration by 14 weeks of age.

**Figure 8.**
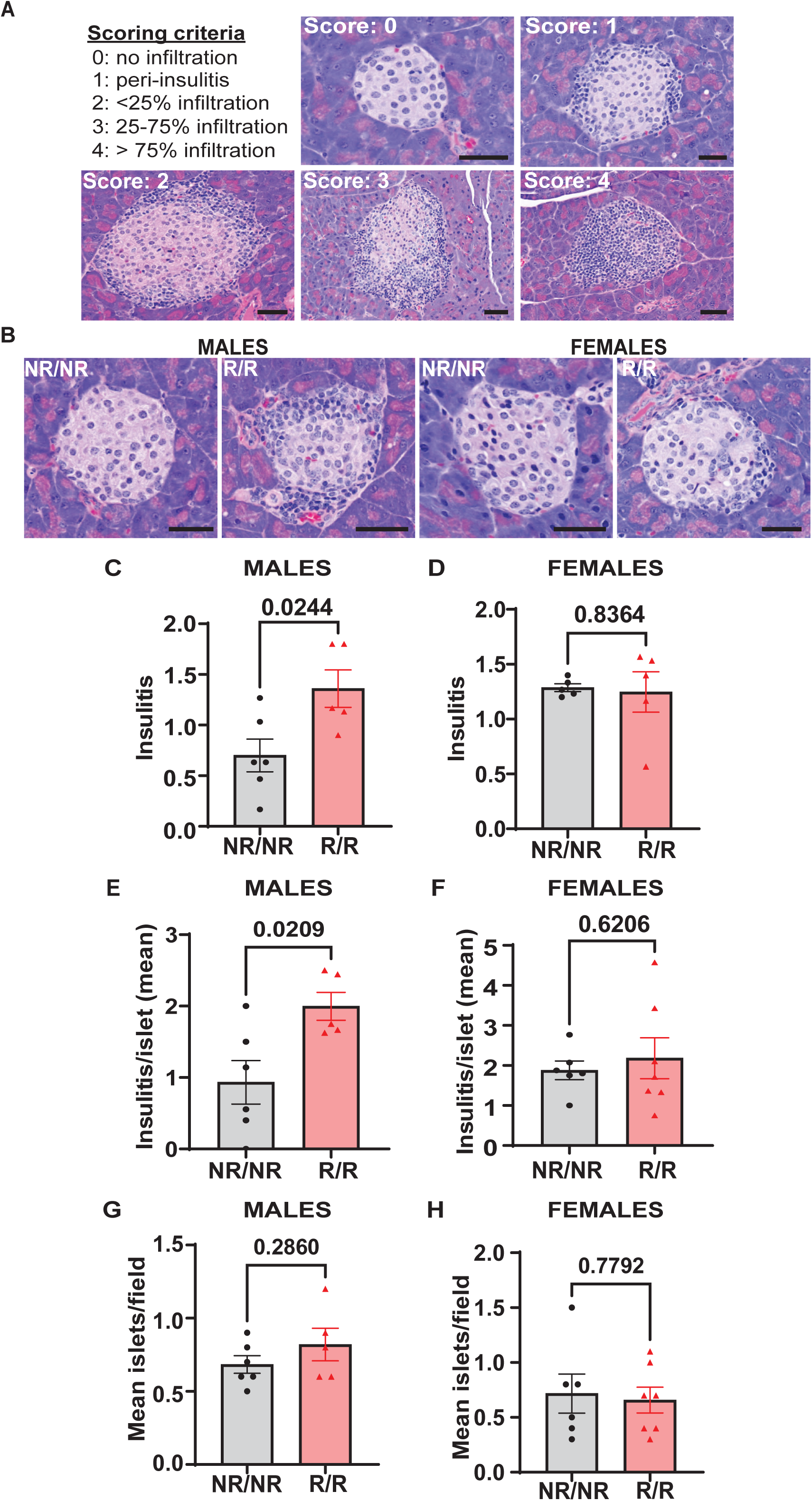
*NOD.Ifih1^R^* increases insulitis in 14-week-old males. **(A)** Representative images of scoring criteria used: 0=no infiltration, 1=peri-insulitis, 2=<25% islet infiltration, 3=25-75% islet infiltration, and 4=>75% islet infiltration. Scale bars: 50μM. **(B)** Representative images showing insulitis from 14-week-old *NOD.Ifih1^NR/NR^* and *NOD.Ifih1^R/R^* mice. Scale bars: 50μM. **(C-D)** Insulitis scores from 14-week-old **(C)** males (*n*=6 *NOD.Ifih1^NR/NR^*, *n*=5 *NOD.Ifih1^R/R^*) and **(D)** females (*n=*5 per genotype). Scores were excluded if less than 10 islets/section could be identified. **(E-F)** insulitis/islet in 14-week-old **(E)** males (*n*=6 *NOD.Ifih1^NR/NR^*, *n*=5 *NOD.Ifih1^R/R^*) and **(F)** females (*n*=6 *NOD.Ifih1^NR/NR^*, *n*=7 *NOD.Ifih1^R/R^*). **(G-H)** mean islets/field in 14-week-old **(G)** males (*n*=6 *NOD.Ifih1^NR/NR^*, *n*=5 *NOD.Ifih1^R/R^*) and **(H)** females (*n*=6 *NOD.Ifih1^NR/NR^*, *n*=7 *NOD.Ifih1^R/R^*), P values were determined using student’s unpaired *t* tests. All data are mean ± SEM.

The increase in insulitis observed in 14-week-old *NOD.Ifih1^R/R^*males prompted us to examine the pancreas for the specific types of immune infiltrates present. We performed flow cytometry to determine the levels of B and T cells in the pancreases. Unlike the spleens, we did not find a significant difference in the frequency of CD8^+^ T cells in the pancreases or draining lymph nodes of *NOD.Ifih1^R/R^* compared to *NOD.Ifih1^NR/NR^*mice **(Fig. S8)**. Additionally, no differences in the frequencies of B cells or plasma cells were detected in the pancreatic draining lymph nodes **(Fig. S8)**. However, a trend for an increased frequency of B cells was observed in the pancreases of 14-week-old males by flow cytometry **(Fig. 9A)**. Therefore, we further investigated B cell infiltration by performing indirect immunofluorescent (IF) staining for B cells on pancreases. Indirect IF for insulin, which is a specific marker for pancreatic β-cells, was also performed to evaluate for β-cell attrition. Strikingly, IF staining revealed a significant decrease in the percentage of β-cell covered area and mean insulin fluorescence intensity per islet as well as a significant increase in the percent of B cell covered area per islet in 14-week-old *NOD.Ifih1^R/R^* compared to *NOD.Ifih1^NR/NR^*males only **(Fig. 9B-E)**. Pancreatic islets from female mice did not show a significant difference in the insulin covered area or fluorescence intensity or B cell covered area **(Fig. 9B, F-H)**. Importantly, the percentage of β-cell covered area in 14-week-old *NOD.Ifih1^NR/NR^* females was comparable to that of 14-week-old *NOD.Ifih1^R/R^* males, and the percentage of B cell covered area in *NOD.Ifih1^NR/NR^* females was even higher than that of *NOD.Ifih1^R/R^* males. These findings further support the notion that the pathology linked to the early stages of diabetes was already more advanced in females compared to males at 14-weeks-old. Furthermore, our results provide evidence that *NOD.Ifih1^R/R^* may promote initial pancreatic B cell infiltration and β-cell death in males.

**Figure 9.**
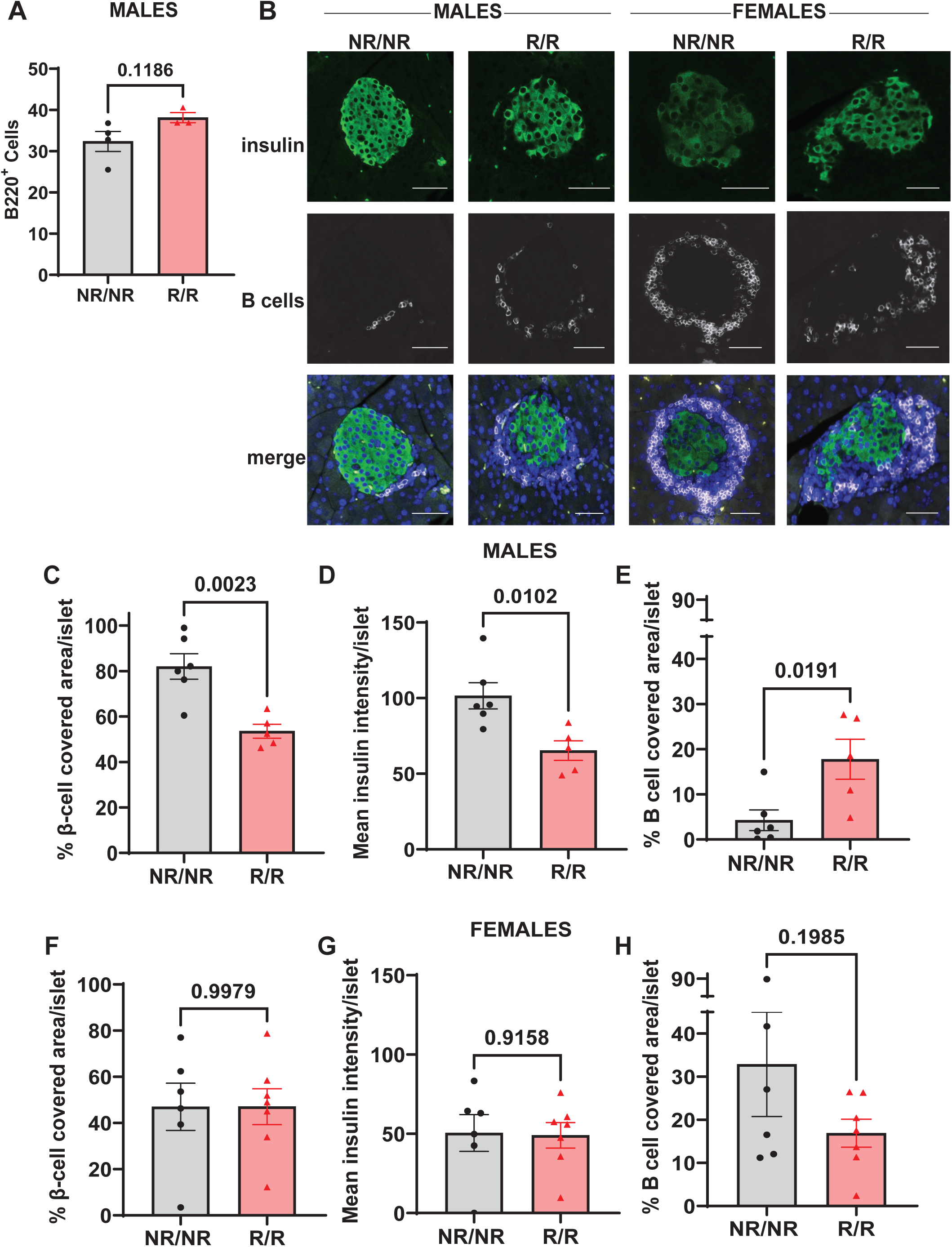
*NOD.Ifih1^R^* promotes depletion of β-cells and enhances B cell infiltration in pancreatic islets in 14-week-old males. **(A)** Frequency of B cells (CD45^+^B220^+^) in the pancreas of 14-week-old male mice (*n*=6 *NOD.Ifih1^NR/NR^*, *n*=5 *NOD.Ifih1^R/R^*) by flow cytometry. **(B)** Representative images showing β-cells (insulin positive, green, top panel), B cells (white, middle panel), and merged images with DAPI (bottom panel). Scale bars: 50μM. **(C-E)** Graphs show **(C)** the percentage of β-cell-covered area/islet, **(D)** mean insulin intensity/islet, and **(E)** the percentage of B cell covered area/islet in pancreases of 14-week-old male mice (*n*=6 *NOD.Ifih1^NR/NR^*, *n*=5 *NOD.Ifih1^R/R^*). **(F-H)** Graphs show **(F)** the percentage of β-cell-covered area/islet, **(G)** mean insulin intensity/islet, and **(H)** the percentage of B cell covered area/islet in pancreases of 14-week-old female mice (*n*=6 *NOD.Ifih1^NR/NR^*, *n*=7 *NOD.Ifih1^R/R^*). P values were determined using student’s unpaired *t* tests. All data are mean ± SEM.

## DISCUSSION

Initiation of T1D pathogenesis is driven by a combination of genetic predisposition(s), environmental risk factors, and anti-viral immunity. Questions remain regarding the precise mechanisms that augment T1D and the interplay among known genetic variants, environmental triggers, and immune-response aberrancies that drive disease onset. Understanding the role of IFIH1^R^ in T1D pathogenesis is important due to its strong propensity to impact disease initiation and/or progression in a multipotent manner. As a cytosolic sensor for viral RNA, IFIH1 is critical for activating IFN I signaling in response to viral pathogens, especially derived from the picornaviridae family which includes CVB, a probable environmental risk factor of T1D. Variants of IFIH1 could alter basal or pathogen-induced cell signaling cascades that influence immune cell populations and susceptibility to autoimmune diseases such as T1D. Since the rs1990760 *IFIH1* risk variant is associated with increased susceptibility to T1D, we set out to determine how IFIH1^R^ alters the immune system by promoting diabetes pathogenesis *in vivo* by utilizing an *Ifih1^R^* knock-in model on the NOD mouse strain. Therefore, we could assess the effect of IFIH1^R^ on diabetes onset and severity. We could also investigate changes indicative of altered interferon signaling and differences in the immune cell populations that might underly the diabetic phenotypes. Our results indicate a causal role of IFIH1^R^ on diabetes development *in vivo*, in which *Ifih1^R^* accelerates diabetes onset in females and alters the frequency and activation of immune cell populations in NOD mice in a sex-dependent and independent manner. In addition, *Ifih1^R^* exhibited an elevated ISG signature, which likely influenced the observed changes in immune cell compartments.

Our previous findings showed that *Ifih1^R^* increased streptozocin (STZ)-induced diabetes incidence as well as synergized with the *Ptpn22* diabetes risk variant (mouse model for the rs2476601 human risk variant) to further accelerate and increase STZ-induced diabetes (21). Our findings in the current study demonstrate that *Ifih1^R^* accelerates diabetes in female NOD mice. As autoreactive cytotoxic CD8^+^ T cells drive pancreatic β-cell destruction in diabetes, it seems plausible that changes in the frequency and activation (CD69^+^ and IFN-γ^+^) of CD8^+^ T cells observed in *NOD.Ifih1^R/R^* mice likely contributed to the accelerated diabetes observed in *NOD.Ifih1^R/R^* females. Supporting an important role for CD69^+^ activated T cells in diabetes, CD69 expression has been found to be increased in T cells from insulin-stimulated whole blood from T1D patients compared to healthy controls (31) as well as in diabetic compared to nondiabetic T cells from NOD mice (32). IFN-γ also plays an important role in T1D pathogenesis by promoting homing of immune cells to pancreatic islets and facilitating β-cell antigen presentation (33, 34). Lack of IFN-γ has been found to delay diabetes development in NOD mice (34). Increased IFN-γ production in memory CD8^+^ T cells likely contributed to accelerated diabetes in *NOD.Ifih1^R/R^*females. Curiously, *NOD.Ifih1^R/R^* lymph nodes displayed similar frequencies of CD8^+^ T cells, but with elevated CD69 expression and frequency of IFN-γ in central and effector memory cells, whereas *NOD.Ifih1^R/R^* spleens showed increased frequencies and numbers of CD8^+^ T cells that displayed decreased levels of CD69. Thus, *NOD.Ifih1^R/R^* may elicit a tissue-dependent effect on CD8^+^ T cells that depends upon additional factors from the surrounding microenvironment. As CD69 has also been considered a retention marker (35), tissue-dependent alterations in CD69 levels could impact the propensity for T cell retention or migration to the pancreas and other sites of immune autoreactivity.

Since we observed an increase in plasma cells in *NOD.Ifih1^R/R^* compared to *NOD.Ifih1^NRN/R^* mice, we posit that increased B cell activation contributed to the accelerated diabetes in females. B cells play a major role in driving T1D by presenting islet antigens to T cells, activating T cells through co-stimulation, and producing islet autoantibodies. Although the onset of T1D can occur at any age, loss of β-cells and disease progression is more rapid in individuals diagnosed at a younger age, and previous studies have attributed this rapid progression to increased pancreatic infiltration and activation of B cells (5, 36). Depletion of B cells by anti-CD20 prevented diabetes in NOD mice (37, 38) and delayed the loss of insulin in T1D in humans, which was more effective in younger patients (5), underscoring the contribution of B cells to T1D pathology. Supporting a role for IFIH1 in diabetes-linked B cell activation, a recent study demonstrated enhanced *Ifih1* gene expression in B cells from a NOD-Rag1^-/-^ somatic cell nuclear transfer (SCNT) B cell receptor model (B1411), which utilized pancreas-infiltrating B cells as donors for SCNT, allowing endurance of B cells in the absence of T cells (39). The B1411 mice showed impaired glucose tolerance and were able to form pancreatic lymph nodes and undergo class-switching in the absence of T cells. Our results showed elevated numbers of plasma cells in *NOD.Ifih1^R/R^* mice, suggesting that *NOD.Ifih1^R/R^* may increase B cell activation. Supporting the notion that elevated plasma cells may have resulted from enhanced IFIH1-mediated downstream signaling, a different *Ifih1* gain of function variant not associated with human disease (G821S) was also found to increase the production of plasma cells in mice (11). Importantly, patients with T1D have shown increased levels of plasmablasts and an increased distribution of plasma cells in peripheral blood samples (40, 41). Thus, we posit that the increased plasma cells present in *NOD.Ifih1^R/R^*may play a causal role in diabetes pathogenesis.

B-T cell interactions enabled by co-stimulatory molecules expressed on the surface of B and T cells are also critical for the formation of plasma cells (42). Concomitant with an increase in plasma cells, our results revealed a trend for increased frequency of B cells expressing CD80 in *NOD.Ifih1^R/R^*mice. CD80 is a co-stimulatory molecule known to drive or inhibit T cell activation by interacting with CD28 or CTLA-4 expressed on T cells, respectively. Moreover, CD80 has also been found to support formation of long-lived plasma cells (27). Increased frequency of CD80 on B cells in *NOD.Ifih1^R/R^* females may have facilitated increased T cell co-stimulation leading to increased plasma cell formation and/or survival. Although plasma cells were increased in *NOD.Ifih1^R/R^* mice, we did not observe differences in autoantibody production. It is possible that differences in IAA may have been more evident in NOD mice at a different/earlier age, as IAA levels have been found to peak between the age 8 and 16 weeks in NOD mice (43). Further, *NOD.Ifih1^R/R^*B cells may be producing other autoantibodies that were not tested. Although an increase in IFIH1 autoantibodies (anti-MDA5 antibodies) have not been reported in T1D, it may be of interest to assess in future studies if *Ifih1^R/R^* impacts development of these autoantibodies, which are deleterious in patients with dermatomyositis (44). Importantly, B cells have the capacity to drive diabetes without producing autoantibodies (45) by acting as antigen presenting cells and co-stimulating T cells. Moreover, plasma cells exert various functions aside from secreting antibodies, including production of cytokines that regulate hematopoiesis and immune tolerance in the intestines (46), which could impact the development of diabetes. Plasma cells or plasmablasts have also been found to be an important source of IL-10 in response to infection or in autoimmunity (28, 47, 48). IL-10 producing plasmablasts/plasma cells were found to be critical for suppressing excessive inflammation in an experimental autoimmune encephalomyelitis mouse model (a mouse model of multiple sclerosis) as well as in mice infected with *Trypanosoma brucei* (28, 47). We speculate that IL-10 producing plasma cells play a regulatory role in the context of T1D. We found that *NOD.Ifih1^R/R^* increased the overall frequency of plasma cells but decreased the frequency of IL-10 within plasma cells of non-draining lymph nodes, suggesting that *NOD.Ifih1^R/R^* may disrupt plasma cell composition or function that could promote inflammation during T1D.

Despite the accelerated onset of diabetes and CD8^+^ T cell activation observed in *NOD.Ifih1^R/R^* female mice, insulitis was more pronounced in *NOD.Ifih1^R/R^* male mice. We speculate that accelerated diabetes coincided with accelerated insulitis in *NOD.Ifih1^R/R^* female mice. Thus, although it is still possible that the age/timing of immune infiltration may differ between *NOD.Ifih1^NR/NR^* and *NOD.Ifih1^R/R^* females, the degree of insulitis was comparable at age 14 weeks. The increased insulitis in 14-week-old males suggests that *Ifih1^R^* can promote pancreatic immune infiltration. This is supported by our findings of increased B cell infiltrates into the pancreas in *NOD.Ifih1^R/R^* male mice. Both B cells and CD8^+^ T cells, with a comparatively higher number of B cells, are known to infiltrate the pancreas in young NOD male and female mice, with infiltration occurring earlier in females (49). The increase in B cells, without an increase in CD8^+^ T cells in *NOD.Ifih1^R/R^* males may also be linked to the timing or stage of disease. Increased B cells may promote enhanced antigen presentation of β-cells that leads to increased frequency or activation of CD8^+^ T cells at a later time point. However, we cannot rule out that regulatory immune cells may have constituted some of the immune infiltrates observed in the islets of *NOD.Ifih1^R/R^* males, which might help explain the more mild diabetic phenotype observed in male compared to female *NOD.Ifih1^R/R^* mice. Therefore, it will be of great interest to further investigate the effect of *Ifih1^R^* on the kinetics and constituents of pancreatic immune cell invasion in NOD mice.

Consistent with the notion that IFIH1^R^ acts as a gain-of-function variant, we observed an increase in the expression of ISGs in *NOD.Ifih1^R/R^* compared to *NOD.Ifih1^NR/NR^* mice, indicative of increased IFN I signaling. As *Mx1*, *Ifit1*, and *Oas1* ISGs were increased in young mice, it seems probable that the increase in these ISGs contributed to diabetes onset. Importantly, an ISG signature is present prior to diabetes onset in children, further supporting that IFN I signaling and increased ISGs may drive disease pathology (7). Since we observed an increase in ISGs in males and females, other sexually dimorphic variables, such as hormones, likely impacted the divergent phenotypes seen in males and females.

Interestingly, a recent study showed that the complete absence of *Ifih1* (*KO*) in NOD mice accelerated T1D in males, which was postulated to occur due to an inability of *Ifih1 KO* mice to produce sufficient myeloid-derived suppressor cells (19). In contrast, female NOD *Ifih1 KO* mice developed disease similarly compared to NOD mice (19). Further, both male and female NOD *Ifih1* mice with an in-frame deletion in the HEL1 domain of IFIH1 (NOD.ΔHel1) exhibited delayed disease onset which was a result of decreased IFIH1-mediated ATP hydrolysis, IFN I production, and pancreatic immune cell infiltration by macrophages, CD4^+^ and CD8^+^ T cells (19). In contrast, we observed accelerated disease in only female *NOD.Ifih1^R^* mice. Despite exhibiting an increase in plasma cells, frequency of splenic CD8^+^ T cells, expression of ISGs, increased β cell depletion, and insulitis marked by increased B cell infiltration at age 14 weeks, *NOD.Ifih1^R/R^* males did not show a significant increase or acceleration in diabetes incidence, suggesting that additional protective mechanisms may have ameliorated disease in males. Taken together, more work is needed to investigate the sex-dependent roles of IFIH1 in T1D pathogenesis.

Collectively, our findings reveal important mechanisms by which IFIH1^R^ may contribute to development of diabetes including upregulation of ISGs, increased insulitis, and immune cell activation. Furthermore, the *NOD.Ifih1* mice present a unique opportunity to investigate how immune-response changes contribute to diabetes. Our results advance knowledge on underlying factors that may connect the *IFIH1* rs1990760 variant with T1D and highlight the need to further investigate the precise role of *IFIH1^R^*in T1D pathogenesis. Follow-up studies should incorporate the use of viral models to determine how the interplay of genetic variants like *IFIH1^R^*and environmental factors, such as CVB, impact diabetes pathogenesis.

## AUTHORS CONTRIBUTIONS

A.J.S. designed and performed experiments, analyzed data and wrote the manuscript. P.G.-P. and L.P.A helped performed experiments and analyzed data. S.D.K. helped analyze data. S.C. and H.B. helped perform experiments and maintain mouse colony. P.W., T.A.Z., A.B.R., E.A., and K.J.M. developed required models/strains and/or helped with writing and editing of the manuscript. J.H.B. and D.J.R. conceived and oversaw the development of the *NOD.Ifih1^A946T^* mice. J.A.G. conceived and supervised the study, analyzed and interpreted data, and helped write and edit the manuscript.

## ACKNOWLEDGMENTS

We sincerely thank Dr. Ben Fowler, Julie Crane, Brie Benjamin-Baker, Rhea Holder, Justin Willige, and Lindsay Stone of the OMRF Imaging Core Facility as well as Jacob Bass and Dr. Diana Hamilton of the OMRF Flow Cytometry core for technical support. The Aurora was supported by NIH S10OD028479-01. We thank Drs. Umesh Deshmukh, Harini Bagavant, Susan Kovats, Jose Alberola-Ila, Joel Guthridge, and Judith James for helpful advice and occasionally providing reagents. We would like to acknowledge Steven Lee for technical support.

## DATA AVAILABILITY STATEMENT

The authors declare that all data supporting this study is available within the manuscript. Additional information is available from the corresponding author upon request.

## Supplementary Material

**Supplementary Figure 1.**
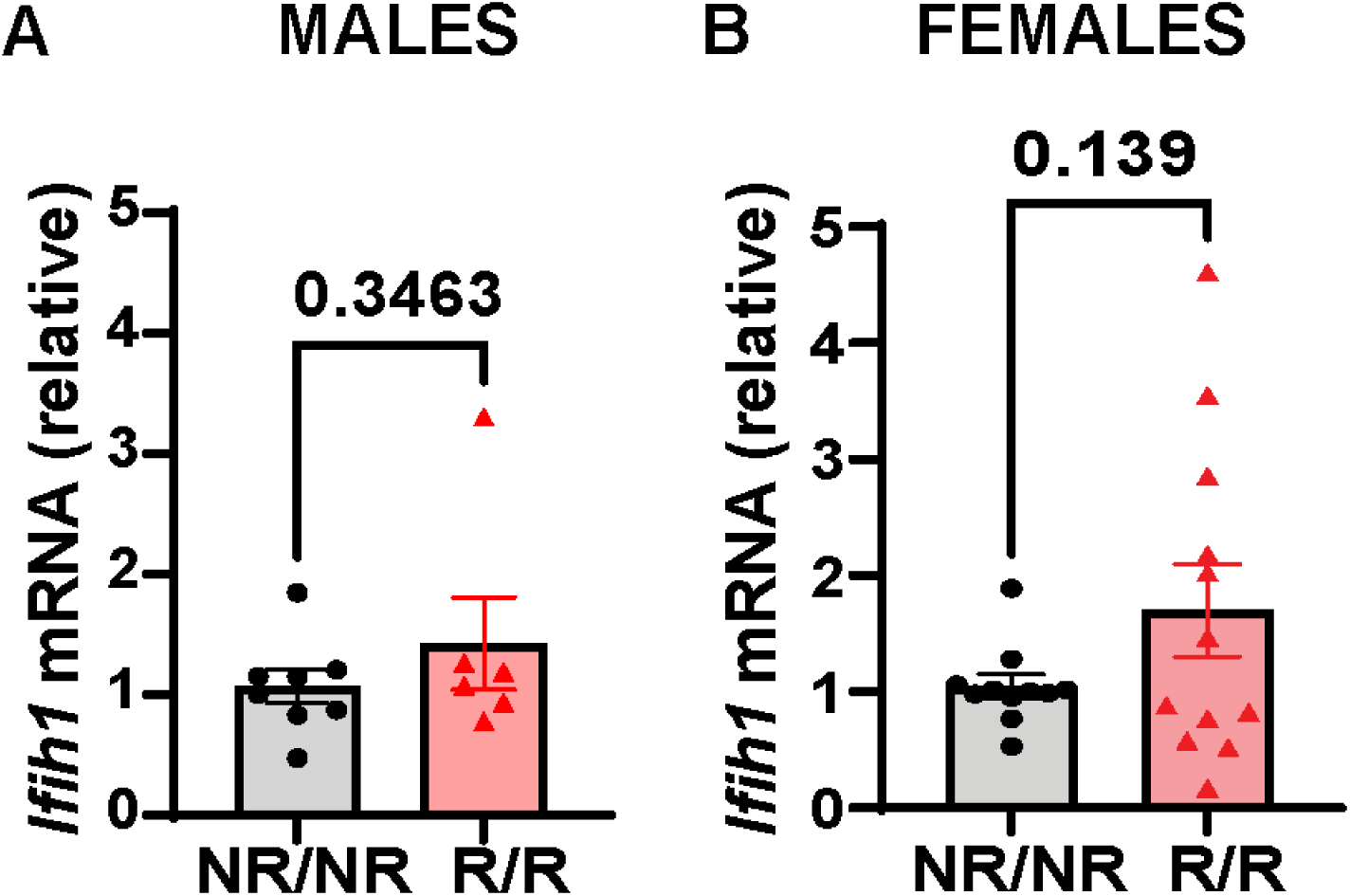
*NOD.Ifih1^R^* does not cause significant changes in *Ifih1* mRNA in 14-week-old mice. The relative mRNA expression of *Ifih1* in 14-week-old **(A)** males (*n*=9 *NOD.Ifih1^NR/NR^*, *n*=6 *NOD.Ifih1^R/R^*) and **(B)** females (*n*=11 *NOD.Ifih1^NR/NR^*, *n*=12 *NOD.Ifih1^R/R^*). P values were determined using student’s unpaired *t* tests. All data are mean ± SEM.

**Supplementary Figure 2.**
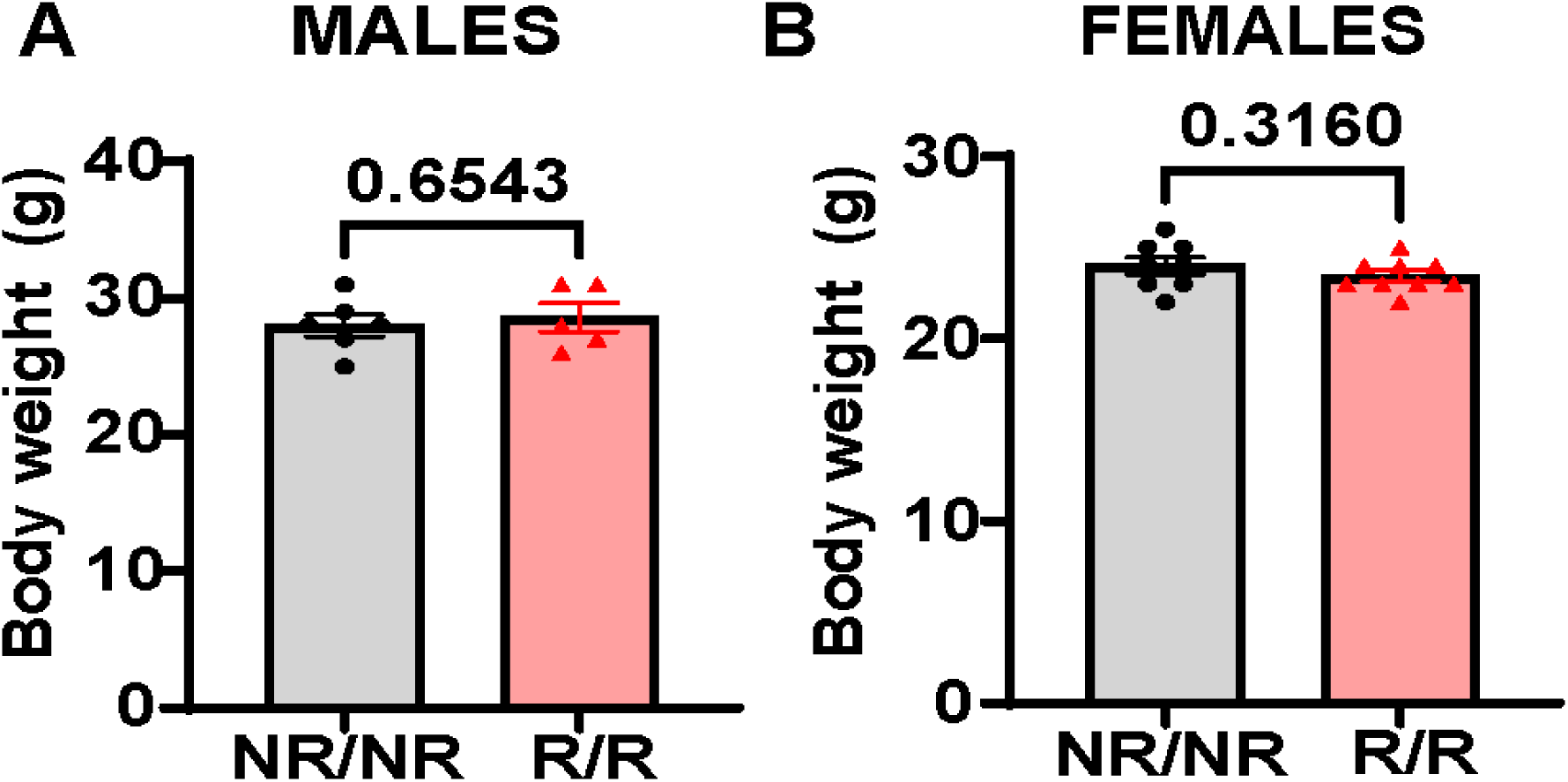
*NOD.Ifih1^R^* does not cause significant changes in body weight. Body weights measured at a single time point immediately prior to euthanasia in 14-week-old *NOD.Ifih1^NR/NR^* compared to *NOD.Ifih1^R/R^* mice in **(A)** males (*n*=6 *NOD.Ifih1^NR/NR^*, *n*=5 *NOD.Ifih1^R/R^*) and **(B)** females (*n*=8 *NOD.Ifih1^NR/NR^*, *n*=9 *NOD.Ifih1^R/R^*). P values were determined using student’s unpaired *t* tests. All data are mean ± SEM.

**Supplementary Figure 3.**
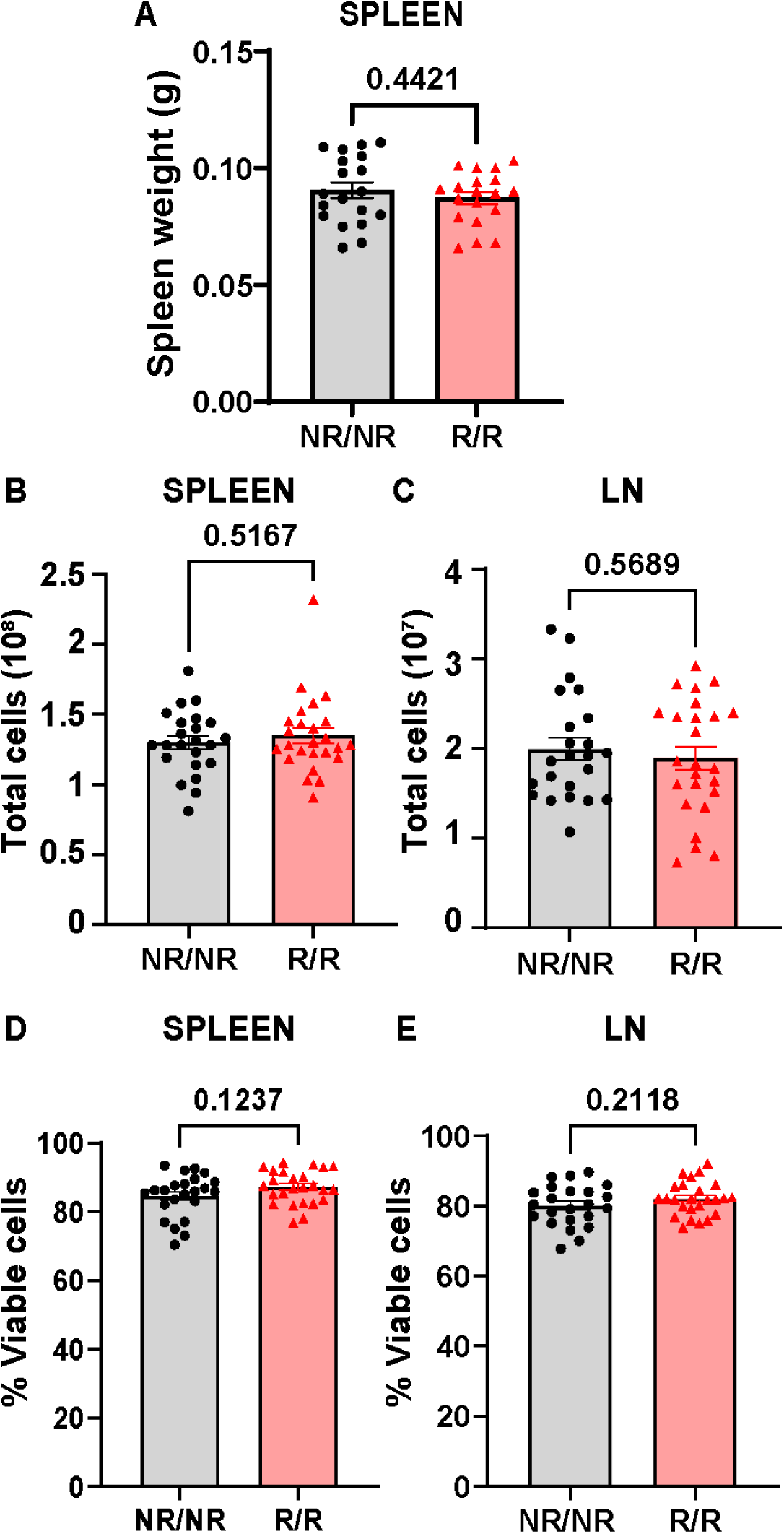
*NOD.Ifih1^R^* does not cause significant changes in splenic weight or cell numbers or viability in spleens or lymph nodes. **(A)** Spleen weight in grams (g) measured at the time of euthanasia in 14-week-old *NOD.Ifih1^NR/NR^* compared to *NOD.Ifih1^R/R^*mice (*n*=19 mice per genotype *NOD.Ifih1^NR/NR^*). **(B-C)** Total number of cells in **(B)** spleens (*n*=23 *NOD.Ifih1^NR/NR^*, *n*=25 *NOD.Ifih1^R/R^*) and **(C)** lymph nodes (LN, *n*=23 *NOD.Ifih1^NR/NR^*, *n*=25 *NOD.Ifih1^R/R^*). **(D-E)** Percent viable cells in **(D)** spleens (*n*=23 *NOD.Ifih1^NR/NR^*, *n*=25 *NOD.Ifih1^R/R^*) and **(E)** lymph nodes (*n*=23 *NOD.Ifih1^NR/NR^*, *n*=25 *NOD.Ifih1^R/R^*). P values were determined using student’s unpaired *t* tests. All data are mean ± SEM.

**Supplementary Figure 4.**
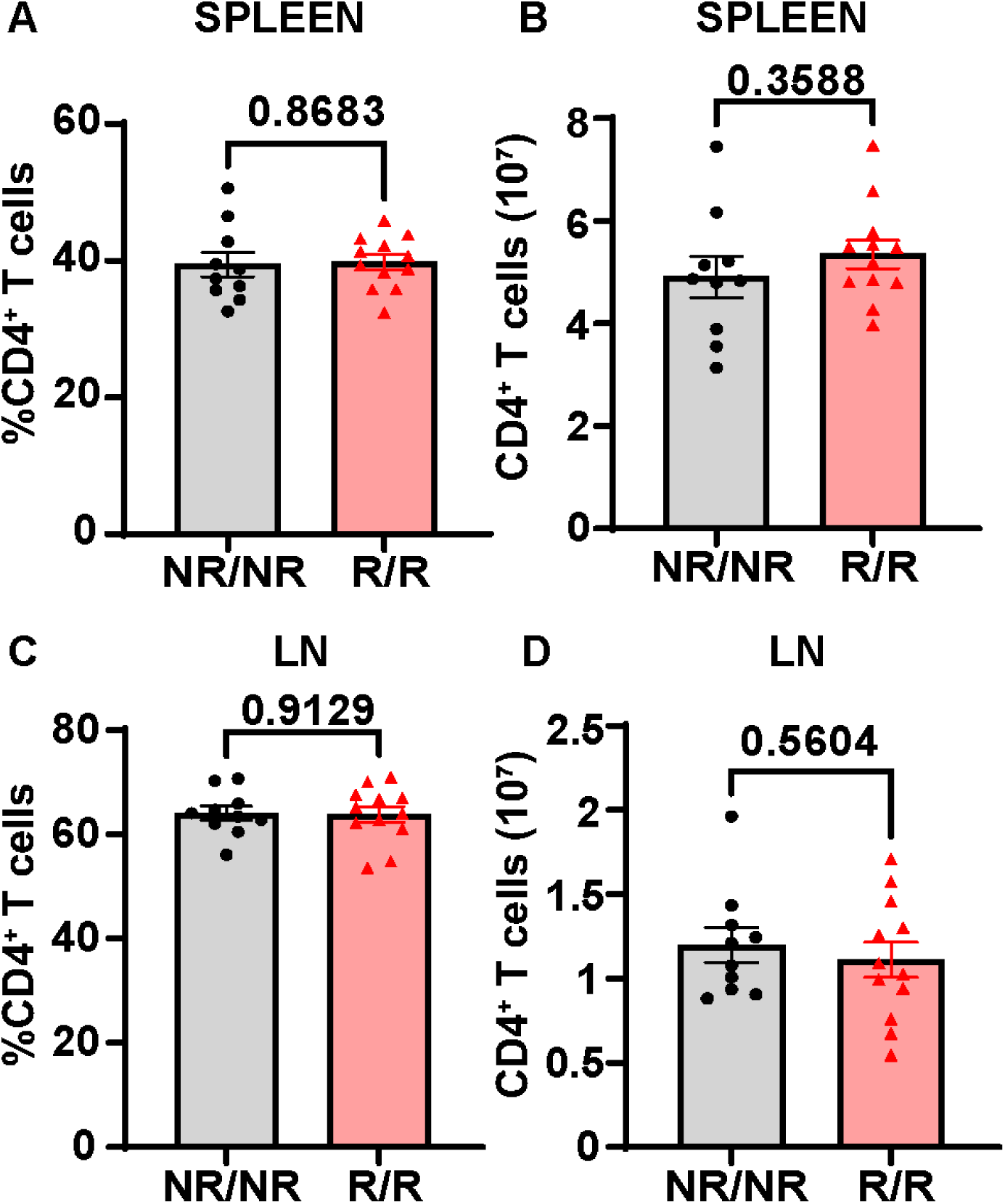
*NOD.Ifih1^R^* does not cause significant changes in the frequency or total cell numbers of CD4^+^ T cells. **(A-B)** CD4^+^ T cells in the spleen as **(A)** frequency of live cells and **(B)** total CD4^+^ cell numbers in *NOD.Ifih1^NR/NR^* compared to *NOD.Ifih1^R/R^*mice (*n*=10 *NOD.Ifih1^NR/NR^*, *n*=12 *NOD.Ifih1^R/R^*). **(C-D)** CD4^+^ T cells in the lymph nodes (LN) as **(C)** frequency of live cells and **(D)** total CD4^+^ cell numbers in *NOD.Ifih1^NR/NR^* compared to *NOD.Ifih1^R/R^* mice (*n*=10 *NOD.Ifih1^NR/NR^*, *n*=12 *NOD.Ifih1^R/R^*). P values were determined using student’s unpaired *t* tests. All data are mean ± SEM.

**Supplementary Figure 5.**
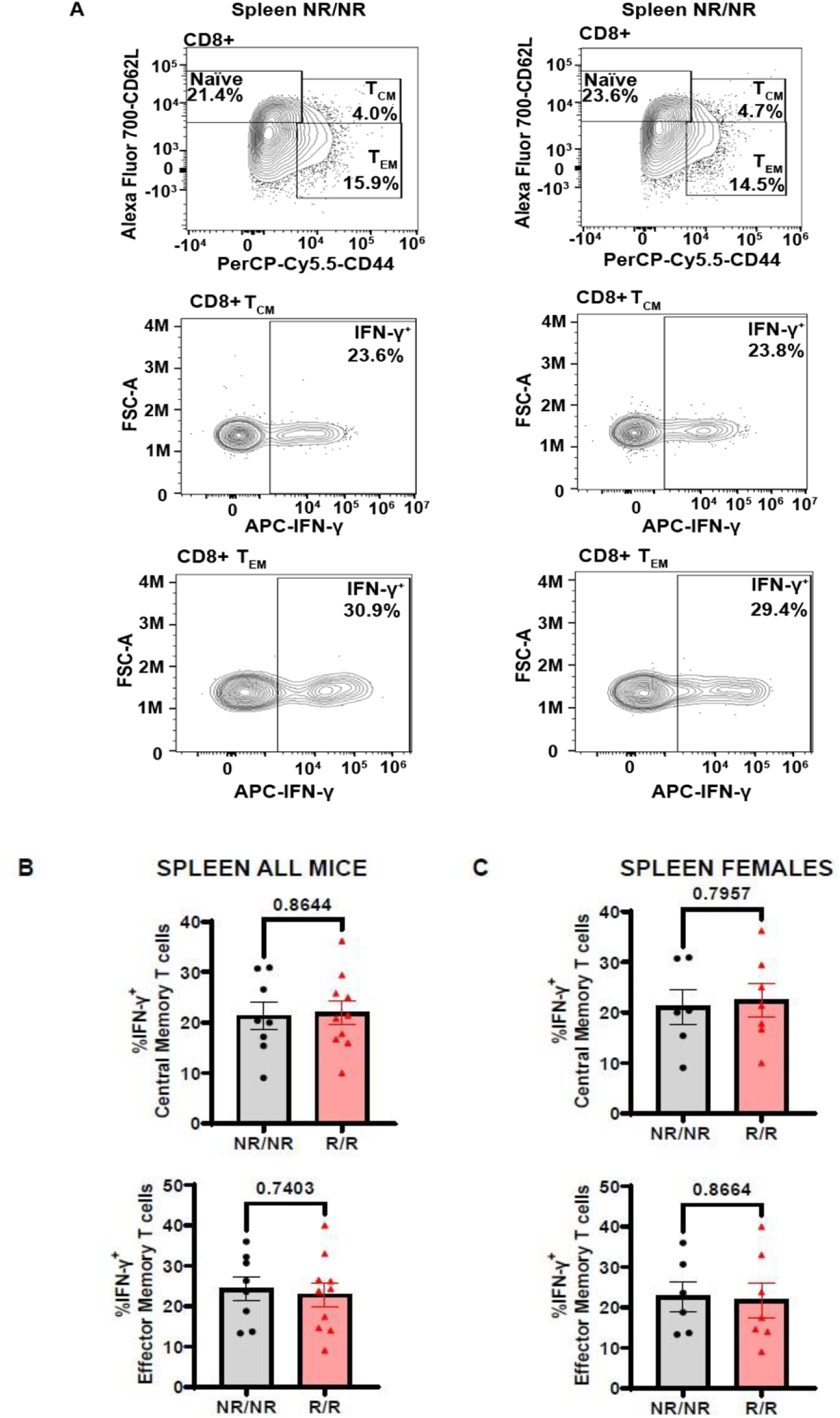
*NOD.Ifih1^R^* does not affect the frequency of IFN-γ on splenic CD8^+^ central memory and effector memory T cells. **(A)** Representative gating of CD8^+^ Naïve (CD62L^+^), central memory (CD62L^+^CD44^+^, TCM), effector memory (CD62L^-^CD44^+^, TEM), and IFN-γ^+^ CD8^+^central memory and effector memory T cells from PMA/ionomycin-treated spleens. **(B-C)** The frequency of IFN-γ^+^ CD8^+^ central memory and effector memory T cells from PMA/ionomycin-treated lymph nodes from **(B)** all mice (*n*=8 *NOD.Ifih1^NR/NR^*, *n*=10 *NOD.Ifih1^R/R^*) and **(C)** females only (*n*=6 *NOD.Ifih1^NR/NR^*, *n*=7 *NOD.Ifih1^R/R^)*. P values were determined using student’s unpaired *t* tests. All data are mean ± SEM.

**Supplementary Figure 6.**
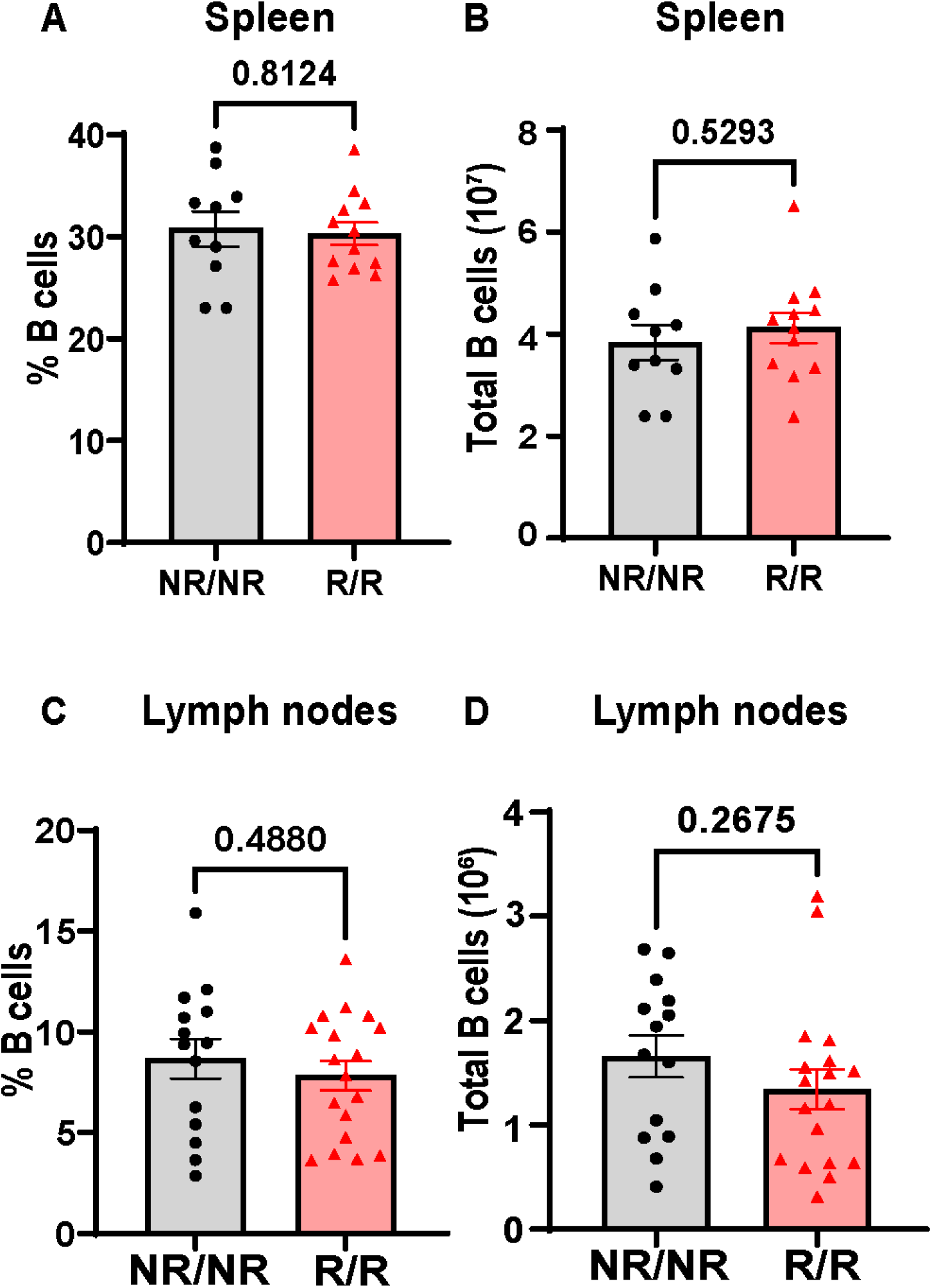
*NOD.Ifih1^R^* does not cause significant changes in frequency or total number of B cells. **(A-B)** Splenic B cells from 14-week-old mice (*n*=10 *NOD.Ifih1^NR/NR^*, *n*=12 *NOD.Ifih1^R/R^*) as **(A)** frequency of live cells and **(B)** total number of splenic B cells. **(C-D)** Lymph node (LN)-derived B cells from 14-week-old mice (*n*=14 *NOD.Ifih1^NR/NR^*, *n*=18 *NOD.Ifih1^R/R^*) as **(C)** frequency of live cells and **(D)** total number of lymph node B cells. P values were determined using student’s unpaired *t* tests. All data are mean ± SEM.

**Supplementary Figure 7.**
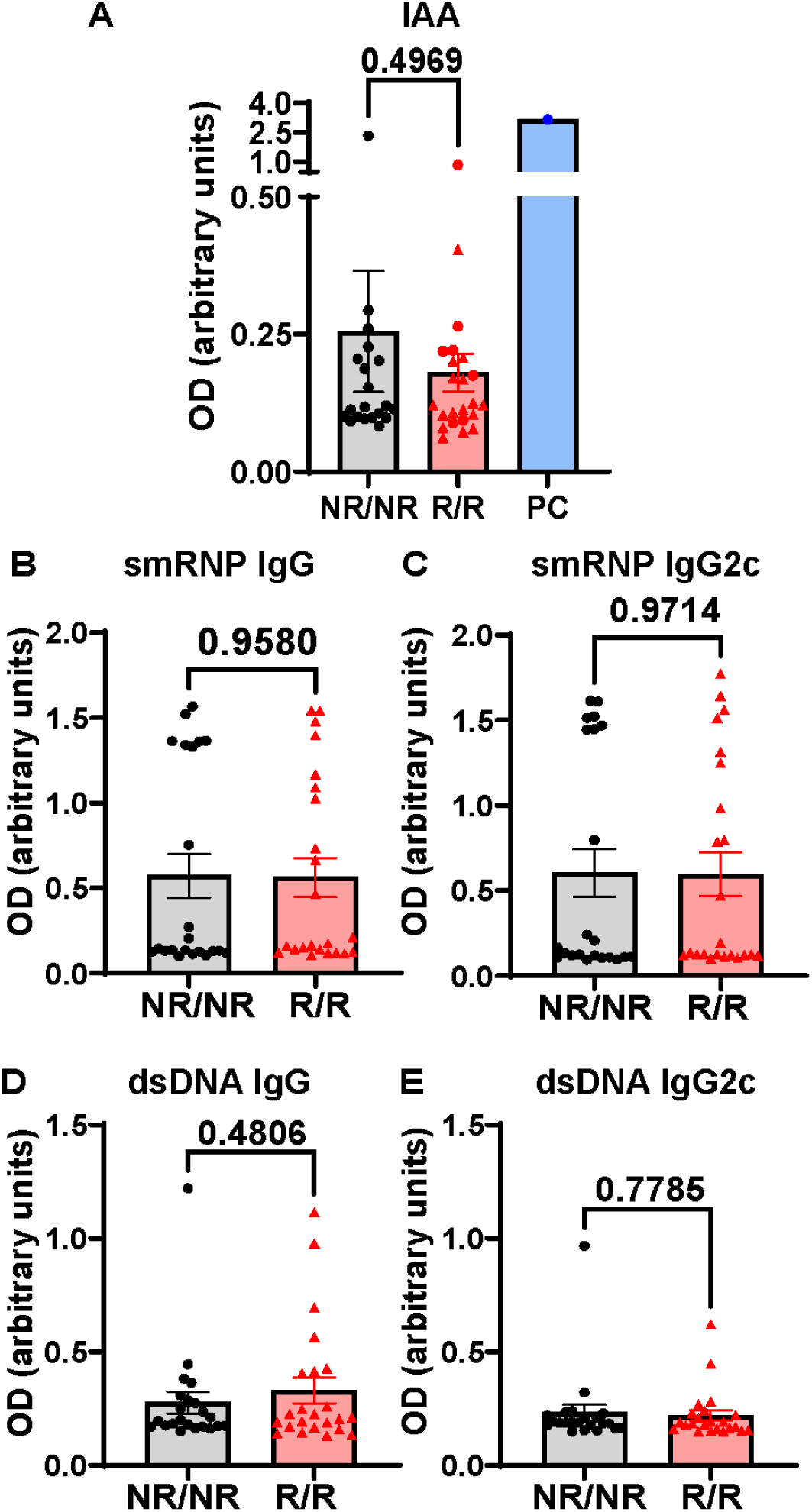
*NOD.Ifih1^R^* does not cause significant changes in insulin, smRNP, or dsDNA autoantibody production. **(A)** OD measurements from Insulin autoantibody (IAA) ELISAs indicative of the levels of IAA in serum from 14-week-old mice (*n*=20 *NOD.Ifih1^NR/NR^*, *n*=23 *NOD.Ifih1^R/R^*). PC: positive control provided by manufacturer. **(B-C)** OD measurements from smRNP **(B)** IgG and **(C)** IgG2c autoantibody ELISAs indicative of the levels of smRNP autoantibody in serum from 14-week-old mice (*n*=22 *NOD.Ifih1^NR/NR^*, *n*=23 *NOD.Ifih1^R/R^*). **(D-E)** OD measurements from dsDNA **(D)** IgG and **(E)** IgG2c autoantibody ELISAs indicative of the levels of dsDNA autoantibody in serum from 14-week-old mice (*n*=22 *NOD.Ifih1^NR/NR^*, *n*=23 *NOD.Ifih1^R/R^*). P values were determined using student’s unpaired *t* tests. All data are mean ± SEM.

**Supplementary Figure 8.**
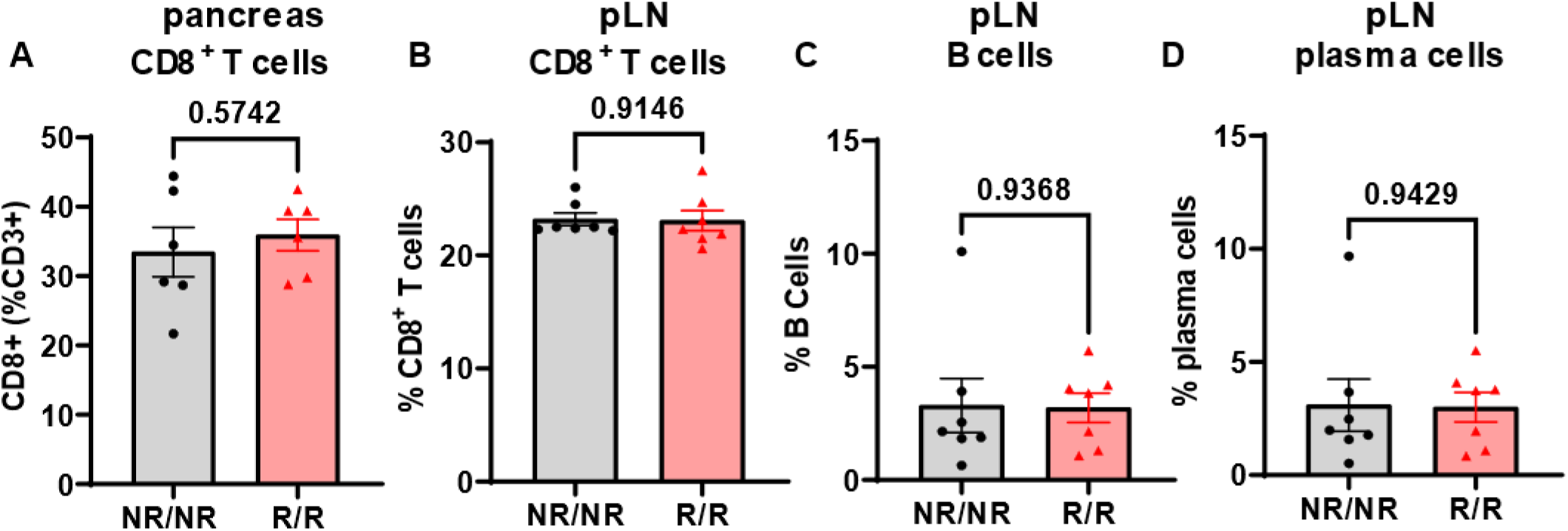
*NOD.Ifih1^R^* does not cause significant changes in the frequency of CD8+ T cells in the pancreas or in the frequency of immune cells in the draining lymph nodes of 14-week-old mice. **(A-B)** The frequency of CD8+ T cells in the **(A)** pancreas (*n*=6 per genotype) and **(B)** pancreatic lymph nodes (pLN, *n*=7 per genotype). **(C)** The frequency of B cells (B220^+^) in the pancreatic lymph nodes (*n*=7 per genotype). **(D)** The frequency of plasma cells (CD19^+^B220^-^ CD138^+^) in the pancreatic lymph nodes (*n*=7 per genotype). All data are from 14-week-old mice. P values were determined using student’s unpaired *t* tests. All data are mean ± SEM.

**Supplementary Table 1.**
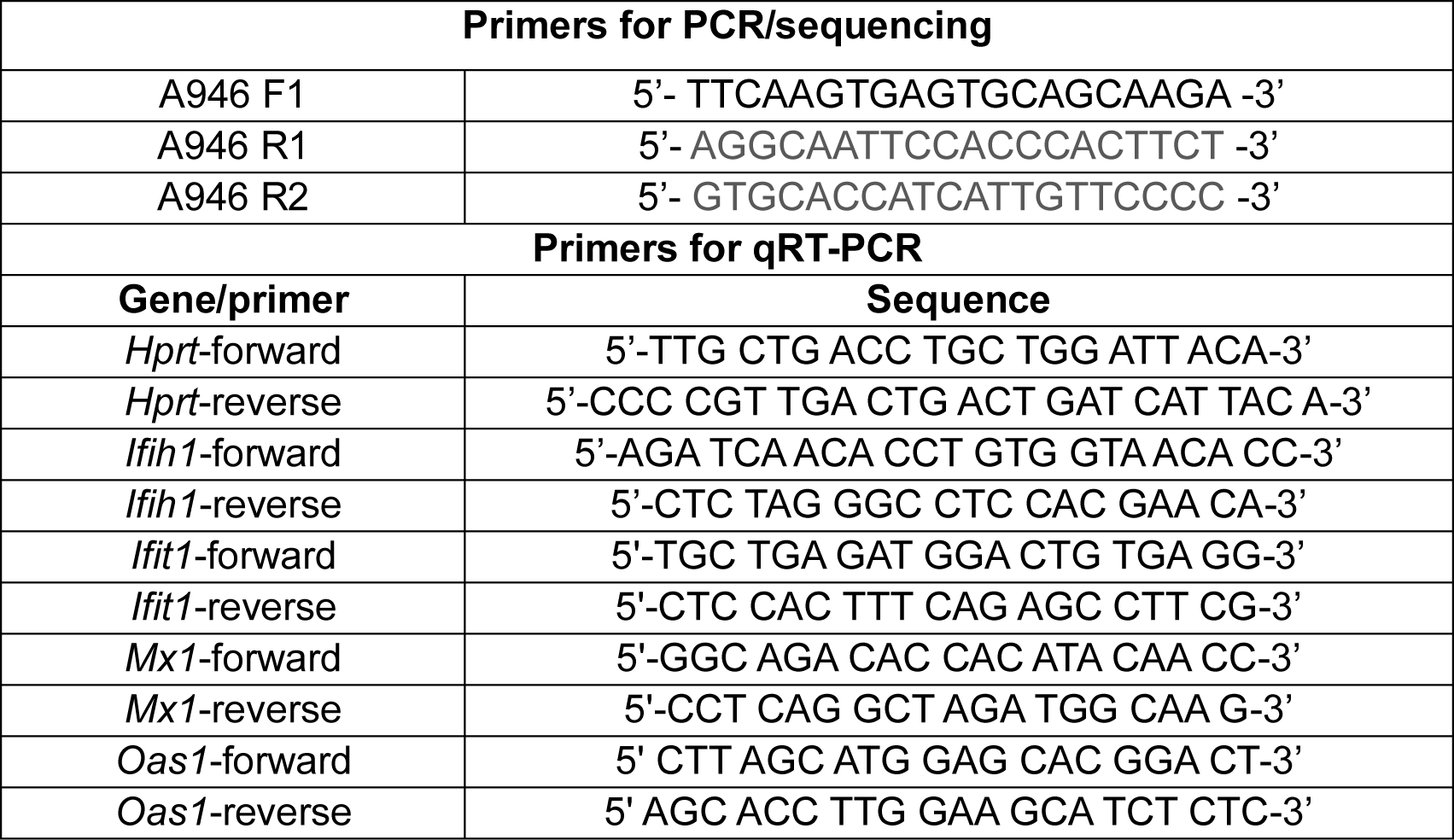
Flow Cytometry Antibodies.

**Supplementary Table 2.** PCR and qRT-PCR Primers.

| **Antibody** | **Company** | **Catalog #** |
| --- | --- | --- |
| BD Horizon™ BUV661 Rat Anti-Mouse CD19 | BD Biosciences | 612971 |
| Brilliant Violet 711™ anti-mouse CD138 (Syndecan-1) | Biolegend | 142519 |
| Near IR fluorescent reactive dye | Thermo Fisher | L34994 A |
| Pacific Blue™ anti-mouse/human CD45R/B220 | Biolegend | 103227 |
| Alexa Fluor® 700 anti-mouse/human CD45R/B220 | Biolegend | 103232 |
| PE anti-mouse CD80 | Biolegend | 104707 |
| PE/Cy7 anti-mouse CD3ε | Biolegend | 100320 |
| PerCP/Cyanine5.5 anti-mouse CD8a | Biolegend | 100734 |
| Brilliant Violet 711™ anti-mouse CD8 | Biolegend | 100759 |
| Spark NIR™ 685 anti-mouse CD69 | Biolegend | 104557 |
| Spark Violet™ 538 anti-mouse CD4 | Biolegend | 100485 |
| APC anti-mouse IFN-γ | Biolegend | 505810 |
| PE-Dazzle 594 anti-mouse IL-10 | Biolegend | 505034 |
| PerCP/Cyanine5.5 anti-mouse/human CD44 | Biolegend | 103032 |
| Alexa Fluor® 700 anti-mouse CD62L | Biolegend | 104426 |
| BD Horizon™ BUV395 Rat Anti-Mouse CD45 | BD Biosciences | 564279 |

